# Genomic vulnerability to climate change and mutation load are affected by past declines in effective population size in two sedentary arctic bird species

**DOI:** 10.1101/2023.01.30.526273

**Authors:** Patrik Rödin-Mörch, Theodore Squires, Kristinn P. Magnússon, Jacob Höglund

## Abstract

Using whole genome re-sequencing data we study the effects of climate influenced declines in effective population size on the accumulation of deleterious mutations and the response to future climate change in populations of cold-adapted avian sister species from the Holarctic: rock ptarmigan (Lagopus muta) and willow ptarmigan (Lagopus lagopus). We reconstruct the demographic histories of the populations and determine their nucleotide diversity, past and present inbreeding, and mutation load. Genomic vulnerability to future climate change scenarios (also known as offset) is predicted for the populations.. We show that relatively small and isolated populations have reduced nucleotide diversity, higher signatures of past and present inbreeding, and higher estimates of mutation load. Among the studied populations, the most vulnerable to a mismatch between current and predicted future environments are rock ptarmigan populations in East Greenland, Iceland, and Svalbard, while among willow ptarmigan, subspecies residing on the British Isles are the most vulnerable.

## Introduction

One of the most imminent threats facing wildlife populations worldwide is climate change (Parmesan 2006; Scheffers et al. 2016). Average temperatures have increased globally and are expected to rise even further in the near future (Collins et al. 2013). The climate is changing at an alarming rate and hence so too are the living conditions of wildlife populations. It is known that climate change has occurred and affected wildlife populations throughout the Earth’s history, as shown during Quaternary periods of glaciation (Hewitt 1996, 1999).Populations and species can respond to climate change in three essential ways: First, species may track the change by altering their range. Second, they may remain in the same place and adapt to the new circumstances via genetic changes, which necessitates that standing genetic variation is available within the population, and/or phenotypic plasticity is present. Third, if neither of the above is possible the populations of such species ultimately go extinct (Parmesan and Yohe 2003, Pinsky et al. 2013, Urban 2015, Höglund et al. 2020).

Since genomes vary in extent of variation both within and among individuals, we do not yet know how much and what kind of genetic variation is needed. In this respect, connectivity among populations plays a crucial role. Various sub-populations of a species can harbor locally adapted genotypes, which may be “pre-adapted” to conditions that will be more common in the future (Lamarque et al. 2013). Thus, it has been argued that dispersal may mitigate the effects of global warming (Aitken et al. 2008), but also that local adaptation may be counteracted by gene flow (outbreeding depression) (Aitken and Whitlock 2013). Alternatively, it is possible that species have become so specialized to a given ecological niche that they cannot adapt to new circumstances especially if environmental change is rapid. A possible recent example is the now-extinct passenger pigeon, (*Ectopistes migratorius*) once claimed to be the most numerous bird species in North America, driven to extinction in the late 19th century by human exploitation. Genomic analyses of preserved specimens have revealed that this bird showed signs of huge past population fluctuations and low genetic variation but also strong signals of selection suggesting that the species had acquired life history characteristics that made it explicitly vulnerable to extinction by humans hunting (Hung et al. 2014, Murray et al. 2017). Thus one of the most pressing scientific issues within contemporary conservation science is: do populations go extinct because of low genetic variation and inbreeding problems, or are they driven to extinction before any loss of genetic variation has impacted them, either by stochastic effects or selection for vulnerable life histories (Höglund 2009, Frankham et al. 2010, Allendorf and Luikart 2012). Here genomic techniques offer a way to study genetic and evolutionary effects of climate change at a fundamental level (Franks and Hoffman 2012, Stillman and Armstrong 2015, Hoffman et al. 2021).

Genetic diversity of a species is shaped by several forces such as mutation, recombination, drift, selection, and gene flow. Among these, selection and gene flow are positively correlated to effective population size, while the effect of drift shows an inverse relationship (Kimura 1983). The impact on genetic diversity during past climatic fluctuations can thus be studied by looking at genome sequence data estimating past changes in effective population size (*N_e_*). Such demographic events determine much of the genetic composition of contemporary populations through their effects on the fundamental processes of mutation accumulation, genetic drift, migration, and selection. Small and isolated populations are predicted to accumulate mutation load (Lynch et al. 1995) and furthermore show higher levels of genetic drift and inbreeding and thereby have reduced genetic variation (Gilpin and Soulé 1986). By their isolation they are also less prone to migration and hence gene flow which may further reduce genetic variation. Finally, the demographic history may have a fundamental influence on the efficacy of selection which is shown theoretically to be much less powerful in shaping allele frequency change in small populations (Kimura 1983). Hence, mutation load may build up and accumulate in small populations (van Oosterhout 2020). However, as levels of inbreeding are also predicted to increase in small populations, deleterious alleles may become exposed to purging which would lead to a reduction of mutation load (Grossen et al. 2020).

On the other hand, deleterious mutations may drift to fixation in small populations (mutational melt down, Lynch et al. 1995). It is therefore useful to discriminate between segregation load (mutations that are still segregating in a diploid population and which may be purged) and drift load (mutations that have become fixed)(van Oosterhout 2020). It is thus not straightforward to predict how the mutation load is affected by demographic events such as bottlenecks (Simons et al. 2014, Mathur and de Woody 2021). All in all, the possibility for natural populations to adapt to future environmental change, and genetic health (susceptibility to inbreeding depression), are predicted to be compromised in small and isolated populations, but there is ambiguity regarding how these factors relate to conservation status (van der Valk et al. 2021, Teixeira and Huber 2021, García-Dorado and Caballero 2021).

In this paper we address these questions by studying two cold adapted and sedentary bird species: the rock ptarmigan (*Lagopus muta*) and willow ptarmigan (*Lagopus lagopus*) (Höglund et al. 2013; Holder et al. 2017). Both are found throughout the Holarctic, however, here we focus on genomic variation in populations on different landmasses surrounding the North Atlantic Ocean. The rock ptarmigan is the more cold adapted of the two species, and mainly occurs in arctic and sub-arctic regions whereas willow ptarmigan, and its subspecies the red grouse (*L. l, scotica*), occur in more open sub-alpine habitats, boreal forests and moorlands (Watson et al. 1998; Storch 2000, 2006; Lucchini et al. 2001). Using whole genome re-sequencing data we determine the population structure and estimate their demographic histories. Furthermore, we determine levels of genetic diversity, past and present inbreeding, and mutation load. Finally, we model genomic vulnerability (offset), defined as the mismatch between current and predicted future genomic variation based on genotype-environment relationships given future climate change scenarios in contemporary populations.

## Methods

### Whole-genome re-sequencing, read mapping and variant calling

Tissue samples from rock ptarmigan were collected from the Pyrenees (France, n=10), the Alps (France, n=8), the eastern (n=10) and the western coast of Greenland (n=9), Iceland (n=10), Sweden (n=6), Svalbard (n=8), and for willow ptarmigan from Newfoundland (Canada n=6), Norway (n=9) and red grouse from northern England (n=9), the latter which technically is considered a sub-species was for analysis purposes grouped with willow ptarmigan. The samples originated from shot birds (wing muscle tissue) with exception of samples from Svalbard (blood samples) which originated from a colony of first-generation captive birds (for detailed sample information including geographical coordinates, see Table. S6).

Most of the DNA was extracted using a high salt extraction precipitation protocol with an extra ethanol precipitation step (modified from Paxton et al. 1996), while Icelandic and eastern Greenland rock ptarmigan DNA was extracted using Monach™ Genomic DNA Purification Kit (New England BioLabs, USA). DNA was checked for concentration and purity using a NanoDrop^®^ 2000 spectrophotometer and Qubit^®^ 3.0 fluorometer Quantitation Kit (Invitrogen™). Libraries were prepared using the TruSeq Nano DNA library preparation kit and sequenced on a S4 flowcell of Illumina Novaseq 6000 by SNP&SEQ technology platform at Uppsala University.

We removed adapters from raw reads using Trimmomatic v.0.36 (Bolger et al. 2014) and evaluated the quality of the reads using FastQC v. 0.11.9 (Andrews 2010). We then mapped trimmed reads onto the chicken (*Gallus gallus*) reference genome (GRCg6a; Genome Reference Consortium; GenBank accession: GCA_000002315.5) using BWA-mem v.0.7.17 (Li and Durbin 2009). We assessed mapping quality and average read coverage using Qualimap v.2.2.1 (García-Alcalde et al. 2012; Okonechnikov et al. 2015), sorted bam files, and marked read duplicates using Picard v.2.20.4 (http://broadinstitute.github.io/picard). In order to decrease variance in sample coverage and facilitate downstream depth filtering across samples, we down-sampled high-coverage samples to ~35X (Table. S6) using samtools v.1.10 (Li et al. 2009). We used GATK v.4.1.1.0 (McKenna et al. 2010) and best practices for variant calling of each species separately (DePristo et al. 2011; Van der Auwera & O’Connor 2020). First, we ran Haplotypecaller with -ERC GVCF enabled on the duplicate marked and down-sampled bam files, followed by combining gVCFs for each individual and chromosome per species using CombineGVCFs, and then jointly genotyped using GenotypeGVCFs. From subsequent vcf with genotyped variants, we separately selected biallelic SNPs and INDELS using SelectVariants. We filtered the resulting VCFs using “hard filtering” standards for the SNP and INDEL variants. First, filtering out SNPs with Phred quality (QUAL) < 30.0, root mean square mapping quality (MQ) < 40.00, strand odds ratio (SOR) > 4.00, variant quality by depth (QD) < 2, fisher strand bias (FS) > 60.00, mapping quality rank sum test (MQRankSum) < −12.5 and read position rank sum test (ReadPosRankSym) < −8.00. For INDELs we used (QUAL) < 30.0, MQ < 40.00, SOR > 10.00, QD < 2.00, FS > 200.00 and ReadPosRankSum < −20.00. We then removed SNPs if they were located within known chicken genome repeat regions as well as if located within 5bp of an identified INDEL using bedtools v. 2.27.1 (Quinlan and Hall 2010). We filtered the final VCF by read depth (15 < DP < 65), allowing a maximum of 10% missing data across samples using vcftools v.0.1.15 (Danecek et al. 2011), resulting in 13,711,844 (rock ptarmigan) and 12,550,543 (willow ptarmigan) variant sites. Furthermore, we assembled mitochondrial genomes from the raw Illumina reads using the Mitofinder pipeline (Allio et al. 2020) utilizing the MEGAHIT assembler (Li et al. 2016) by mapping the raw reads onto known avian mitochondria reference sequences. We removed three individuals with highly fragmented assemblies (Table. S6) and aligned the remaining individuals using Clustal Omega (Sievers et al. 2011). We manually trimmed large gaps in the alignment found in multiple individuals using PhyDE v.0.9971 (http://www.phyde.de/), resulting in a final alignment of 13406bp with 462 segregating sites across all populations and species.

### Phylogenetic relationships and population structure

We merged the VCF containing biallelic SNPs for both species using bedtools v. 2.27.1 (Quinlan and Hall 2010). Using SNPhylo v.20180901 (Lee et al. 2014), we constructed a maximum likelihood phylogenetic tree based on the SNP data with 1000 bootstrap replicates, setting variant pruning based on linkage disequilibrium (LD>0.2) with a maximum window size of 500kb and minor allele frequency (MAF < 0.1) resulting in 3,205,910 variant sites. Using FitgTree v. 1.4.4 (http://tree.bio.ed.ac.uk/software/figtree/) the tree was plotted as a cladogram with midpoint rooting. We constructed a median joining network from the mito-genome alignments using PopART (http://popart.otago.ac.nz). Using ADMIXTURE v. 1.3.3 (Alexander et al. 2009) we estimated ancestry coefficients using the same filtered vcf used in the SNPhylo analysis, by conducting separate runs for each species (*L.muta:* K2-7, *L.lagopus:* K2-5) and plotted the results using CLUMPAK (Kopelman et al. 2015).

### Genomic diversity and offset under future climate change scenarios

We calculated autosomal nucleotide diversity (π) using pixy v.0.95.0 (Korunes and Samuk 2021), which provides an unbiased estimate of nucleotide diversity by including invariant sites in its calculation. To include invariant sites in our vcf we re-ran GenotypeGVCFs in GATK with the setting “--include-non-variant-sites”. Furthermore, we calculated nucleotide diversity for the mitogenome using Arlequin v.3.5.2 (Excoffier and Lischer 2010). In order to assess rock and willow ptarmigan vulnerability to future climate change we estimated the genetic offset (here referred to as genomic vulnerability to be consistent with previous literature) between allele frequency associated with the current environment and associations with the environment projected under four different climate changes. We estimated offset using LEA v.3.3.2 (Gain and François 2021) which represent the miss-match due to genetic drift between a population with allele frequencies associated with current climate variation and a hypothetical version of that population with allele frequencies associated with future climate variation, where the offset statistic has the same interpretation as FST. This method assumes weak and polygenic effects of the environment spread throughout the genome. We filtered the combined vcf allowing no missing data and a maf of 5% and converted the vcf to plink format using vcftools v.0.1.15 (Danecek et al. 2011). In order to obtain a better model fit, we LD pruned the data in Plink v.1.9 (Chang et al. 2015; www.cog-genomics.org/plink/1.9/) based on slightly less stringent criteria than for the construction of the phylogenetic tree, trimming one variant in a variant pair with r^2^ > 0.5 in windows of 50 SNPs with a 5 SNPs step size, resulting in a total of 385,676 variants across populations and species. The genetic offset method implemented in LEA requires an initial gene-environment analysis (GEA) using latent factor mixed models (LFMM) on current environmental variables. In order to estimate the optimal number of clusters we calculated ancestry coefficients using the sparse non-negative matrix factorization (snmf) function in LEA and its cross-entropy criterion for each K number of ancestral populations (Fig. S1). We performed GEA using the new implementation of LFMM with ridge regularization that estimates per locus effect sizes from multivariate environmental data. Even though the optimal number of ancestral populations is 6, we assumed 9 latent factors corresponding to the number of sampled populations, in order to account for the large geographical separation among all populations. For each sampling location we utilized 18 current climate variables from the WorldClim database (https://worldclimg.org), having to drop one due to perfect collinearity with other variables, and projections from the four Representative Concentration Pathway (RCP) trajectories for the year 2070 (RCP 2.6, RCP 4.5, RCP 6.0 and RCP 8.5) which all differ in areas such as projected emissions, global land use and population size (IPCC 2014).

### Demographic history

We estimated effective population size (N_e_) over two different time spans using two different methods. To estimate more recent trends in N_e_, we used the linkage disequilibrium (LD) based method implemented in SNeP v.1.1 (Barbato et al. 2015). This method estimates N_e_ in different time periods by dividing the genome into bins based on the distance between biallelic SNPs and averaging LD over bins, where bins of larger distance are more informative about recent changes and short distance about N_e_ further back in time. To estimate N_e_ from physical distance the minimum and maximum recombination rate (cM) is inferred by utilizing one of the predefined mapping functions available in the software. We split the vcf based on population origin and converted it to ped an map files using PLINK v.1.9 (Chang et al. 2015; www.cog-genomics.org/plink/1.9/). We ran SNeP with the following settings; a minimum of maximum distance between SNPs of 50kb and 4 Mb, minor allele frequency cut-off of 0.1, a maximum of 100 000 SNPs per chromosome, a bin-width of 50kb and corrections for limited sample size and unphased data. Since we do not have an inferred genome-wide recombination rate, we assumed the default standard recombination rate (10^-8^) to convert physical distance into cM used by the mapping function to infer the recombination rate between pairs of SNPs. To estimate changes in N_e_ in a more distant past, we used pairwise sequentially markovian coalescent (PSMC v.0.6.5; Li and Durbin 2011) that infers N_e_ at different time periods for a single autosomal diploid genome, based on the distribution of time to the most recent common ancestor for independent genomic segments and the rate of coalescent events. We generated the fastq file containing the diploid consensus sequence using mpileup in samtools together with bcftools view and vcfutils v.0.1.18 (Li 2011) with adjust mapping quality (-C) set to 50, filtering based on read depth lower than 12 and higher than 70 and base quality below 20. We ran PSMC with 34 free atomic time intervals (4+30*2+4+6+10) for a maximum of 30 iterations, with a maximum 2N_0_ coalescent time and initial theta/rho ratio set to 5. In order to estimate the uncertainty of our estimate we performed 100 bootstrap replicates by sub-sampling random genomic stretches of 5Mb from the diploid consensus sequence.

### Runs of homozygosity (ROH) and relative mutation load

The length distribution of runs of homozygosity (ROH) is due to recombination that is informative about the processes shaping these segments and their relative age, where length classes between 100kb – 1Mb (short ROH) represent genetic drift resulting from historic demographic changes and long stretches in the range of > 1 Mb represent more immediate inbreeding (Ceballos et al. 2018). Using PLINK v.1.9 (Chang et al. 2015; www.cog-genomics.org/plink/1.9/). We estimated runs of homozygosity (ROH) for each population by setting the minimum length to 100Kb, the maximum gap between two consecutive SNPs to be part of the same ROH to 1000kb and the minimum SNP count for a ROH to 50. Furthermore, we characterized the population level relative mutation load using the Ensembl variant effect predictor tool (VEP, McLaren et al. 2016) with sorting intolerant from tolerant activated (SIFT, Ng and Heinkoff 2003), running the software for each individual separately in order to identify and predict the influence of synonymous, missense, deleterious, and loss-of-function mutations (transcript ablation, splice acceptor variant, splice donor variant, stop gained, frame-shift variant, stop lost, and start lost) based on the chicken genome and annotations. We conducted separate gene ontology enrichment analyses per population using g:Profiler (Raudvere et al. 2019), and analyzed the genes located within short and long ROH, missense, deleterious and loss-of-function mutations separately using the native algorithm to correct for multiple testing employing a genome-wide threshold 0.05.

### Predictors of mutation load

To estimate which predictor or combination of predictors, without interactions, best explain observed patterns of population-level variation in missense, deleterious and loss-of-function mutations, we fitted a series of multiple regression models using in R v.3.6.3 (R Core Team 2021) and conducted AICc based model selection using the AICcmodavg package (Mazerolle 2020). We constructed our models using a combination of the following parameters; the harmonic mean of recent changes in effect population size estimated with SNeP and historic changes inferred with PSMC, as well as short and long ROH and latitude (Table. S5).

## Results

### Phylogenetic relationships and population structure

The maximum likelihood phylogenetic tree reveals an expected split between rock ptarmigan and willow ptarmigan. The sub-species red grouse from the British Isles clustered together with Norwegian willow ptarmigan and Canadian willow ptarmigan were found to be more divergent (Fig. S2). There are two main lineages within rock ptarmigan, one consisting of Greenland and Iceland in the west, and another containing mainland European populations from Sweden and France (the Pyrenees and the Alps) with the population from Svalbard forming a sister group to the remaining rock ptarmigan populations. The ADMIXTURE results show that for willow ptarmigan at K2 (Fig 1a), red grouse and Norwegian willow ptarmigan cluster together, while at K3 (Fig 1a) there is more substantial influence from English red grouse within the Norwegian willow ptarmigan cluster than from individuals from Newfoundland. The cross validation error estimates used to assess the optimal K reveals a steady increase with increasing K and thus an optimal K cannot be detected for willow ptarmigan (Fig. S3a). For rock ptarmigan at the optimal value of K (K=4, Fig. S3c), the populations cluster in correspondence to the phylogenetic tree (Fig. S2), where Greenland and Iceland cluster together separated by a cluster containing mainland European populations from the French Alps and Sweden, and where the French Pyrenees and Svalbard form their own clusters (Fig. 1b). For K equal to the number of sample populations (K=7) each population forms their own cluster without much admixture,however, the population from the western coast of Greenland reveals substantial sub-structure.

**Fig.1.**
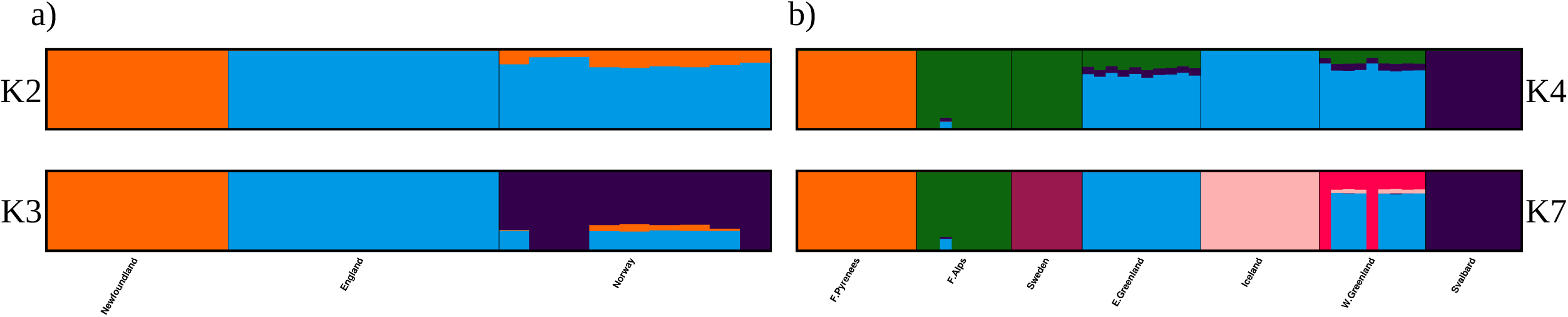
Ancestry coefficients estimated by Admixture, (a)For number of clusters K4 and K7 for rock ptarmigan, (c) K2 and K3 for willow ptarmigan.

The mitochondrial medium joining network reveals high haplotype diversity within the three willow ptarmigan populations and a clear separation from Rock ptarmigan (Fig. S4). Within rock ptarmigan we find more variation in the number of haplotypes among populations but isolated populations such as those in the French Pyrenees and Svalbard only contain two haplotypes. Furthermore, the shared ancestry between Iceland and Greenland and the influence from Svalbard indicated by the ADMIXTURE analysis appears to be reflected in the absence of any clear haplotype structure among two populations from Greenland and Iceland, while Svalbard groups fairly closely to all three populations.

### Genomic diversity and offset under future climate change scenarios

We find that autosomal nucleotide diversity (π) ranges between 3.6*10^-3^ – 4.3*10^-3^ in willow ptarmigan and for mitochondria 8.7*10^-3^ – 3.3*10^-2^ (Table. S1). In rock ptarmigan autosomal π ranges from 9.3*10^-4^ – 3.2*10^-3^ and 3*10^-5^ – 4.9*10^-2^ for mitochondria, while the lowest nucleotide diversity can be found in the populations from Svalbard (autosomal: 9.3*10^-4,^, mitochondria: 3*10^-5^), the French Pyrenees (autosomal: 1.5*10^-3^, mitochondria: 3*10^-5^), and Iceland (autosomal: 1.7*10^-3^, mitochondria: 2.7*10^-2^).

In order to predict the future genomic response to climate change (genomic vulnerability) we use the estimate of genetic offset between current and predicted future environment, whose interpretation is analogous to F_ST_ and where an estimated offset close to 1 is considered extremely high. We find that the genetic offset is between 0.077 – 0.981 for rock ptarmigan and 0.194 – 0.789 for willow ptarmigan (RCP2.6, Fig. 2a), 0 – 0.981 and 0.186 – 0.988 (RCP4.5, Fig. 2b), 0.33 – 0.987 and 0.065 – 0.912 (RCP6.0, Fig. 2c), and finally 0 – 0.983 and 0.171 – 0.951 for rock ptarmigan and willow ptarmigan respectively (RCP8.5, Fig. 2d). Most populations of both species are fairly exposed to future climate change, which would indicate that they might not have enough time to adapt to rapidly changing conditions; this is especially the case for the populations located close to or above the Arctic Circle. The most vulnerable population to climate change under all four future projections was Svalbard for rock ptarmigan (genetic offset 0.981 – 0.987) and English red grouse for willow ptarmigan (genetic offset 0.789 – 0.951).

**Fig.2.**
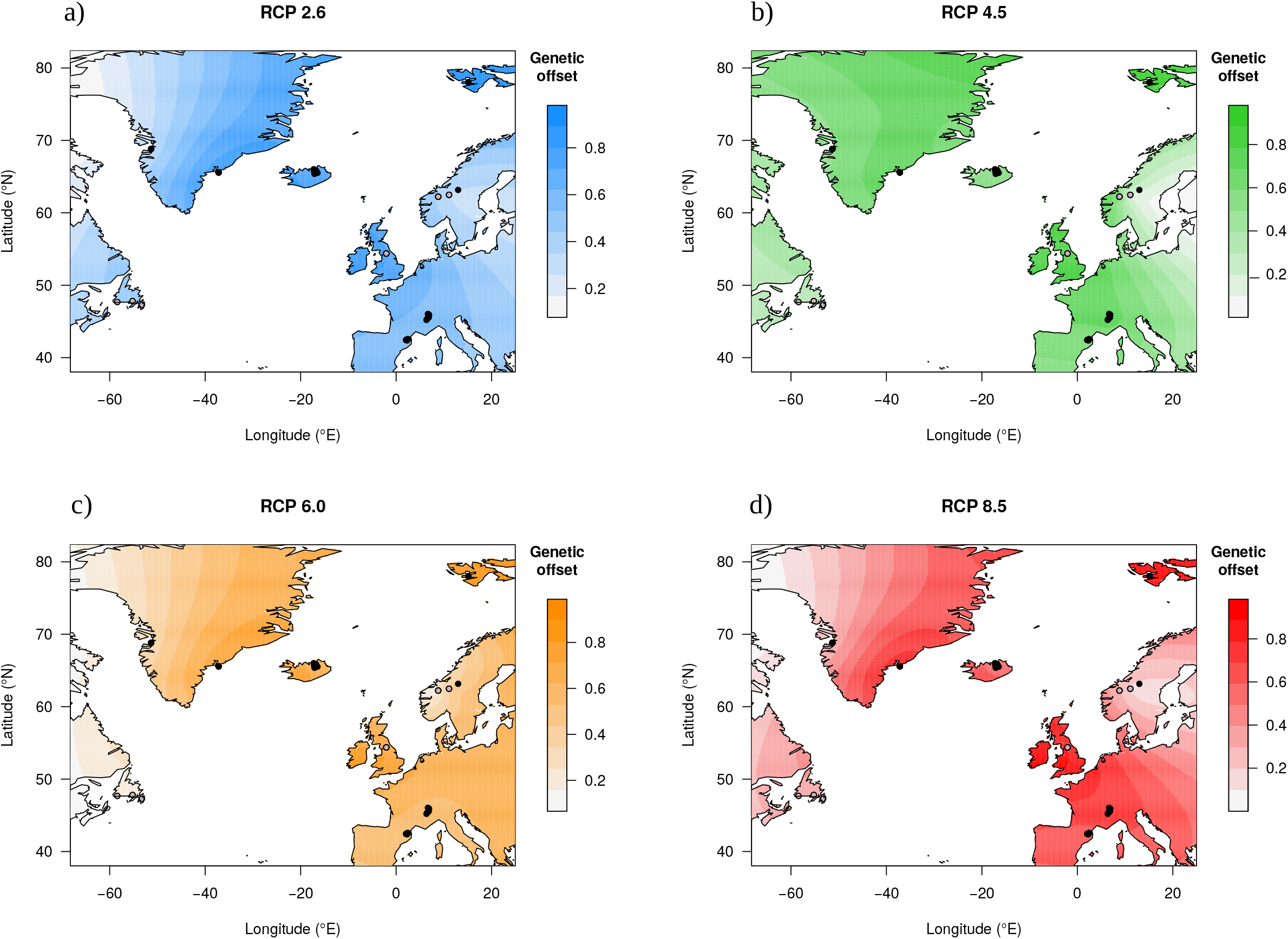
Genomic vulnerability estimated as genetic offset to future climate change under four future climate change scenarios for rock ptarmigan (black points) and willow ptarmigan (grey points). Higher values means larger offset between current and future climate, where the latter consists of the representative concentration pathways (RCP) describing four scenarios of green house gas emissions of varying severity, (a) RCP2.6, (b) RCP4.5, (c) RCP6.0, (d) RCP8.5.

### Demographic history

The estimates of contemporary trends of effective population size using SNeP, reveals approximately 2000 years of declining N_e_ for both rock and willow ptarmigan (Fig. 3a,c). Rock ptarmigan started out with a N_e_ ranging between 1500-3100, where isolated populations such as those from Svalbard and the French Pyrenees were on the lower end, followed by a steady decline until present day where the range of N_e_ is considerably lower (N_e_ = 22 – 47). Estimated willow ptarmigan N_e_ 2000 years ago ranged between 1500-3400, followed by a similar decline as rock ptarmigan until present day (N_e_=22 – 43). For historic estimates of N_e_ using PSMC we find quite large differences between populations and species but with similar trajectories (Fig. 3b,d; Fig. S5a-j). The overall trend of N_e_ for rock ptarmigan starting some 10^6^ years before present (YBP) until roughly 20,000 YBP, was that of a population increase starting 10^6^ YBP that reached a maximum N_e_ between 80,000 – 200,000 YBP, where we find the largest N_e_ in the French Alps, and the smallest estimates of N_e_ in populations with small current N_e_ (i.e. Svalbard, the French Pyrenees, and Iceland). Since, all seven populations showed a substantial decrease in N_e_ for the remainder of the recoverable demographic history. We observe the same pattern for willow ptarmigan, but where peak N_e_ was reached further back in time compared to most rock ptarmigan populations (200,000 – 250,000 YBP), together with an overall higher maximum N_e_. For willow ptarmigan we find the largest N_e_ in the Norwegian population and where English red grouse and the population from Newfoundland show roughly similar N_e_.

**Fig.3.**
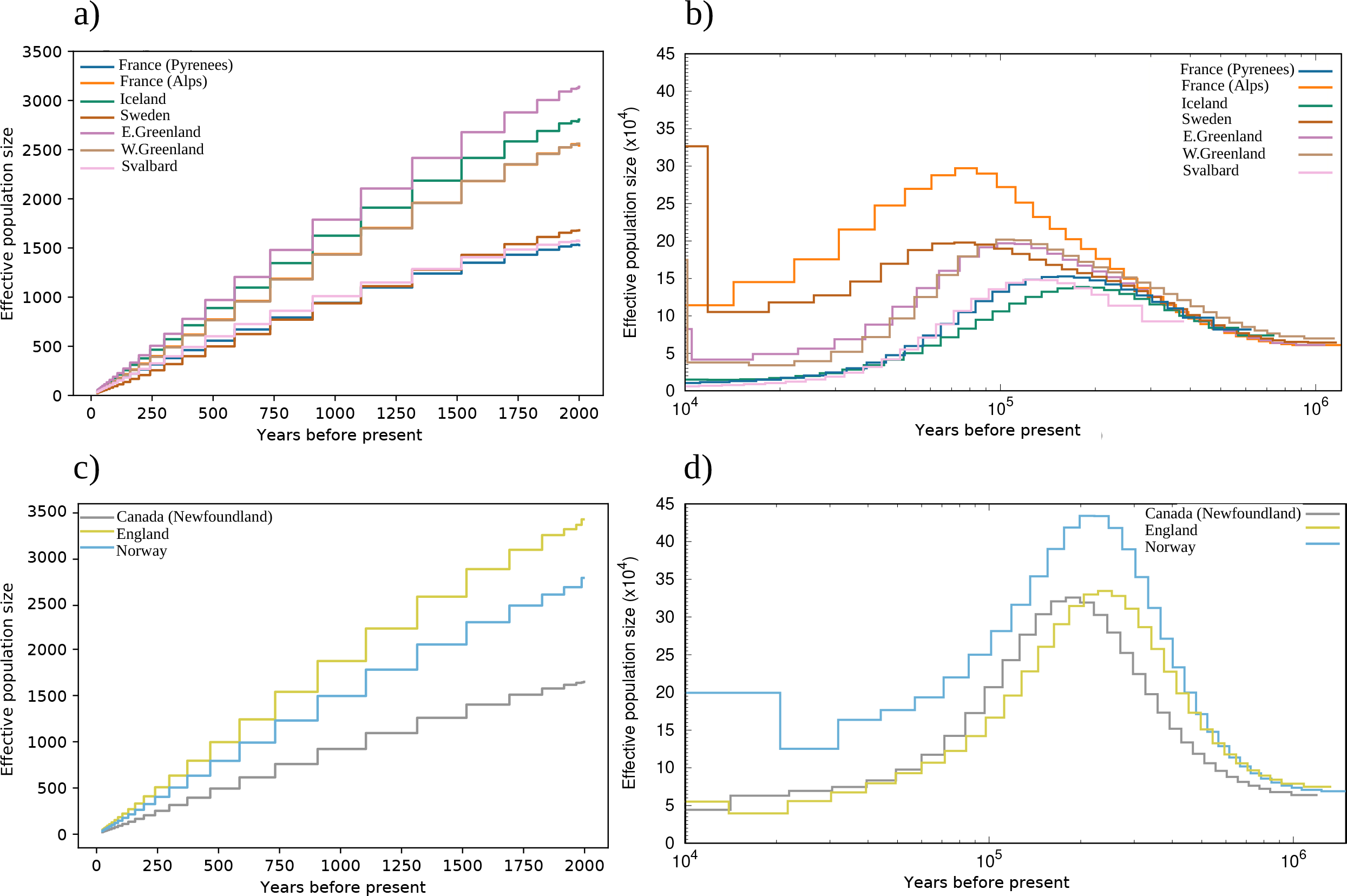
(a) Recent trends in effective population size (N_e_) using SNeP for rock ptarmigan, (b) historic N_e_ trends estimated using PSMC for rock ptarmigan, (c) recent N_e_ trends for willow ptarmigan, (d) historic N_e_ trends for willow ptarmigan.

### Runs of homozygosity (ROH) and relative mutation load

The percentage of the autosomal genome covered by runs of homozygosity (ROH) of rock ptarmigan consists on average of 0.04 – 3.4% long ROH and 1.4 – 18.4% short ROH where we find the higher percentages in populations with smaller current and historic N_e_ (Table. S1). Both cumulative length (Mb) of short and long ROH differ among population (short: F_9,75_ = 204.4, p<0.0001, long: F_9,75_ = 32.63, p<0.0001, Fig. S6). Most rock ptarmigan populations show comparable lengths of long ROH, but where the genome of individuals from populations from the French Pyrenees and Svalbard consists of longer stretches and larger number of ROH than the other populations (Fig. 4b, Fig. S6). For willow ptarmigans we find that English red grouse and willow ptarmigans from Norway have less of the genome represented as long ROH compared to Newfoundland. For willow ptarmigan, the cumulative length of short ROH is higher in the Newfoundland and English red grouse populations compared to Norwegian willow ptarmigan.

**Fig.4.**
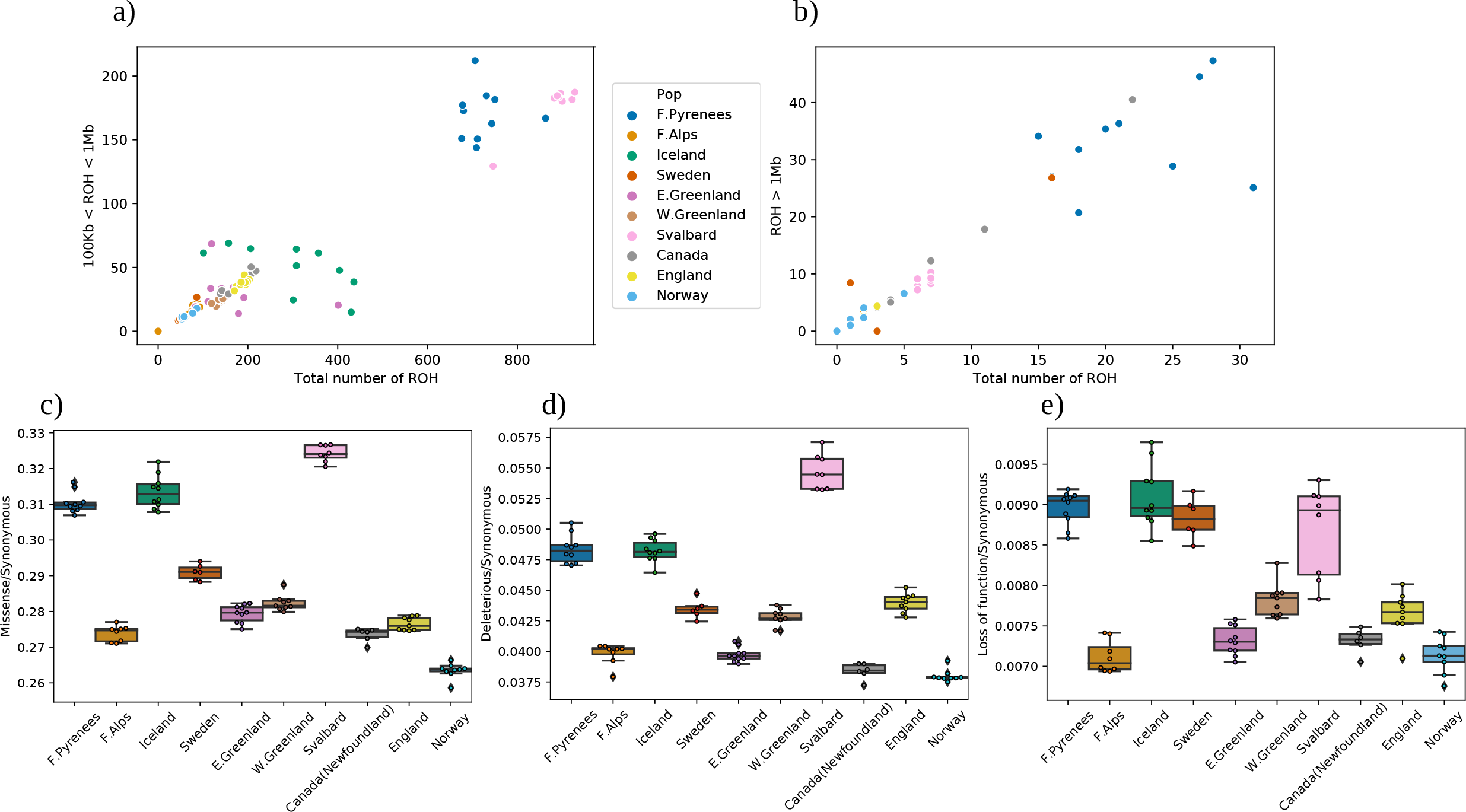
(a) Cumulative length of short ROH (Mb) plotted against the total number of short ROH, (b) Cumulative length of long ROH (Mb) plotted against the total number of long ROH, relative mutation load (c) missense mutations, (d) deleterious mutations, (e) loss-of-function mutations.

All three categories of relative mutation load considered here differ among populations (missense: F_9,75_ = 502.9, p<0.0001, deleterious: F_9,75_ = 304.5, p<0.0001, loss-of-function: F_9,75_ = 67.6, p<0.0001). We generally find higher relative mutation load in rock ptarmigan populations than willow ptarmigan reflecting the differences in maximum N_e_ between the two species (Fig. 4c,d,e, Fig. S7a,b,c). The highest relative mutation load based on the fraction of missense and predicted deleterious mutations to synonymous mutations for rock ptarmigan can be found in Svalbard, Iceland and the French Pyrenees. In willow ptarmigan, these measures are higher in the populations from Newfoundland and English red grouse compared to Norwegian willow ptarmigan. The fraction of loss of function to synonymous mutations shows similar patterns among populations compared to missense and deleterious mutations with the rock ptarmigan populations from Svalbard, Iceland, and the French Pyrenees showing slightly higher within species levels. Within willow ptarmigan we observe more even levels of loss-of-function to synonymous, but with slightly elevated levels in the Swedish population and English red grouse. A large proportion of each individual class of mutation segregates in the populations are fairly low frequencies (Fig. 74a,b,c).

We do however find that mutations within the rock ptarmigan populations from the French Pyrenees, Svalbard and Iceland segregates at higher frequencies with more fixed deleterious alleles (Table. S2a,b,c; Fig. S7a,b,c: missense median allele frequency = 0.4-0.6, IQR = 0.5-0.7; deleterious median allele frequency = 0.2-0.5, IQR = 0.6-0.625; loss-of-function median allele frequency = 0.5-0.7, IQR = 0.5-0.6) compared to the other populations (missense median allele frequency = 0.222-0.333, IQR = 0.5-0.667; deleterious median allele frequency = 0.125-0.333, IQR = 0.25-0.556; loss-of-function median allele frequency = 0.25-0.444, IQR = 0.5-0.667). Examples of the most significantly enriched GO terms (Table. S3) for missense mutations involves biological processes such as double-strand break and DNA repair, for deleterious mutations; cillium organization and assembly and no significantly enriched terms for biological processes for loss-of-function mutations. Only a relatively small number of all mutations that we classify here as contributing to the relative mutation load overlaps with ROH (Table. S4) ranging from 0-11.1% (missense mutations), 0-10.1% (deleterious mutations) and 0-8.4% (loss of function mutations), where, as we would expect, the populations containing the highest fraction of relative load shows the largest overlap. Most overlaps between mutation and ROH are with short ROH owing to the fact that all of the populations has much fewer long ROH relative to short ones. This might however, indicate less effective purging in the populations with the highest overlap between the three classes of mutation load and ROH.

### Predictors of mutation load

Based on model selection of a number of multiple regression models (Table. S5) we find that variation in relative load based on missense mutations is best explained by short ROH and harmonic mean historic N_e_ estimated by PSMC (R^2^_adjusted_= 0.838, F_2,82_ = 218, p < 0.0001), the same combination of predictors also explains variation in relative load of deleterious mutations (R^2^_adjusted_= 0.796, F_2,82_ = 164.9, p < 0.0001). For loss-of-function mutations however, we find that a model containing harmonic mean of recent and historic N_e_ estimated by SNeP and PSMC respectively, is the best fitting model (R^2^_adjusted_= 0.572, F_2,82_ = 57.22, p < 0.0001).

## Discussion

In this study we find in two sedentary and cold adapted birds species large differences in genetic variation among the geographically separated populations, with declining effective population sizes through time, which has resulted in higher levels of inbreeding and mutation load in isolated island populations in the North Atlantic as well as alarming patterns of genomic vulnerability to future climate change.

### Phylogenetic relationships and population structure

Both species in this study have either truly circumpolar (rock ptarmigan) or near circumpolar (willow ptarmigan/red grouse) distributions (Storch 2000), where local populations are connected by gene flow creating patterns of isolation by distance (Sahlman et al. 2009, Höglund et al. 2013). This explains why Newfoundland willow ptarmigan are more distant from Norwegian willow ptarmigan than British red grouse: the North Atlantic Ocean acting as an effective barrier to gene flow. In the case of rock ptarmigan, the North Atlantic Ocean has historically been a less obvious barrier as this species is adapted to even more extreme arctic climates and have hence been able to colonize Greenland and islands in the north like Iceland and Svalbard from which willow ptarmigan are absent. Previous studies have suggested that Svalbard has been colonized by rock ptarmigan from the east (Taymyr Peninsula), while the origin of birds colonizing Iceland has been more obscure (Sahlman et al. 2019). Here we find evidence of influence on the Icelandic population mainly from Greenland (autosomal data) in the west and weak but possible influence from Svalbard in the east (autosomes and the mitogenome).

### Demographic history

We find clear evidence of a relation between past and present population size, demographic history and levels of isolation on genetic diversity in both species. First, willow ptarmigan populations tend to display higher overall levels of genetic diversity than rock ptarmigan in line with a hypothesis of more connected populations allowing higher gene flow in the former species. Rock ptarmigan are adapted to more extreme arctic conditions and are thus found at higher elevations and further north than willow ptarmigan. Hence, this more cold adapted of the two species has been able to remain at high elevations at southern latitudes even after past warming of the climate, such as in the Pyrenees and the French Alps and has been able to colonize islands, such as Iceland and Svalbard, further north than willow ptarmigan. This has had the consequence that rock ptarmigan populations are relatively isolated with historically smaller population sizes.

However, PSMC analyses of both species show population decline starting about 200,000 years ago in willow ptarmigan and somewhat later in most rock ptarmigan populations except from Scandinavia and the Alps. These latter two populations would be the ones that remained in contact for the longest with the main distribution on the Palearctic landmass (Lagerholm et al. 2017). Furthermore, all sampled populations show consistent declines in recent times. These more recent declines are possibly also an effect of anthropogenic influences like habitat destruction and hunting.

### Inbreeding and mutation load

The past demography also shed light on the patterns of inbreeding levels and mutation load. The relatively more isolated rock ptarmigan populations were generally shown to be more affected by past (as estimated by short ROH) and more contemporary inbreeding levels (long ROH). It is interesting that the Svalbardic population of rock ptarmigan together with those from the French Pyrenees show the most extreme levels of short ROH, consistent with historical inbreeding in line with persistent small population size and isolation. While Svalbardic birds are not extreme when it comes to long ROH, those from the Pyrenees remain on the extreme side. This indicates that Svalbard was either colonized from the Eurasian landmass (Sahlman et al. 2009) quite recently or the rock ptarmigan population has been influenced by gene flow in recent times while the Pyrenees have been isolated for longer.

Not surprisingly, missense mutations are much more common than mutations classified as deleterious or causing loss-of-function, the latter category beig the most uncommon. In line with patterns observed for genetic diversity and ROH, willow ptarmigan populations tend to differ in terms of mutation load from rock ptarmigan. Willow ptarmigan populations lie in the lower end when comparing populations across the two species, while some rock ptarmigan populations show higher levels of mutation load. Again, the isolated populations in the Pyrenees, Iceland, and Svalbard stick out as relatively extreme. We would again attribute this pattern to the effect of the past demography where purifying selection removing deleterious alleles is predicted to be less efficient in small isolated populations (Kimura 1983). Furthermore, in such populations deleterious alleles may drift towards fixation (mutational melt down, Lynch and Gabriel 1990). Both effects likely apply in the case of the populations studied here. This scenario is strengthened by our observation that variation in relative load based on both missense and deleterious mutations was best explained by short ROH and harmonic mean historic N_e_ estimated by PSMC. For loss-of-function mutations, however, we find that a model containing harmonic mean of recent and historic N_e_ estimated by SNeP and PSMC, respectively, was the best fitting model, indicating that loss-of-function mutations may be more affected by contemporary events resulting in more immediate effects of inbreeding.

### Genomic diversity and offset under future climate change scenarios

Previous studies integrating population genomics and environmental data have identified genomic variation associated with climate across the breeding range of a migratory songbird, the yellow warbler (*Setophaga petechial*, Bay et al. 2018). Warbler populations requiring the greatest shifts in allele frequencies to keep pace with future climate change had experienced the largest population declines, suggesting that failure to adapt may have negatively affected populations. Furthermore, climate factors and socioeconomic parameters were used to predict genomic vulnerability in a Central African rainforest songbird, the little greenbul (*Andropadus virens*, Smith et al. 2020). Similarly, genomic vulnerability has been used to predict future change and adaptation in plants (e.g. Dauphin et al. 2020, Rhoné et al. 2020).

The factors discussed above have all combined to affect the genomic vulnerability of grouse populations around the North Atlantic. Genomic vulnerability is here defined as the mismatch between current and predicted future genomic variation based on genotype-environment relationships modeled across contemporary populations (Gain and François 2021). The effect on any ptarmigan population appears more dependent on geographical location than any modeled climate scenario or even species identity. In rock ptarmigan, East Greenland, Iceland, and Svalbard populations are predicted to become most impacted while populations in West Greenland and Scandinavia are projected to be the least affected. In willow ptarmigan, British red grouse are predicted to be most affected while Newfoundland willow ptarmigan are predicted to become least affected. These predictions are based on a combination of the predicted local climate offset and the standing genetic variation in the contemporary populations. The fact that northern populations are identified among the most vulnerable is line with observations of Arctic ecosystems being the most impacted by climate change (Box et al. 2019).

Future model predictions of climate responses hold great promise but are not unchallenged. Incorporating genomic data in such modeling efforts may improve forecasts as this allows for populations to adapt to the predicted environmental changes (Hoffmann et al. 2021). In a study of two cryptic European forests bats it was concluded that by considering climate-adaptive potential, range loss predictions were reduced as compared to scenarios which did not allow for adaptive evolution (Razgour et al. 2019). Our study shows that populations on islands in the far north of the Atlantic Ocean are predicted to be the most vulnerable. This is especially worrisome since Arctic ecosystems are projected to be the ones most affected by rising temperatures (Box et al. 2019). Furthermore, species with limited dispersal capacity, such as ptarmigans may not easily change their range by moving. Further, for the more cold-adapted rock ptarmigan, there is limited habitat available to open further north for some populations like those in Svalbard.

## Data Availability

Upon acceptance of the manuscript, raw sequencing data will be uploaded on SRA. Output files will be uploaded on dryad and analysis scripts will be made available on dryad and PRMs GitHub page.

## Acknowledgments

We would like to thank all people who have contributed with samples to make this study possible. In particular Nicolas Bech, Jean-Francoois Allienne, Claude Novoa, Jerome Bossier, David Newborn, Chris Callohar, Daniel Appenroth and Ólafur Karl Nielsen. We would would also like to thank Theodore Squires for valuable input on the manuscript.

The sequencing was performed by the SNP&SEQ Technology Platform in Uppsala. This facility is part of the National Genomics Infrastructure (NGI) Sweden and Science for Life laboratory. The SNP&SEQ Platform is supported by the Swedish Research Council and the Knut and Alice Wallenberg Foundation. Uppsala Multidisciplinary Center for Advanced Computational Science provided computing resources. This study was financially supported by the Swedish Research Council (project 2018-04635 to JH) and Icelandic Research Fund (Project 206529-051 to KPM).

