## Supplementary_material for "Genomic vulnerability to climate change and mutation load are affected by past declines in effective population size in two sedentary arctic bird species"

### Supporting information

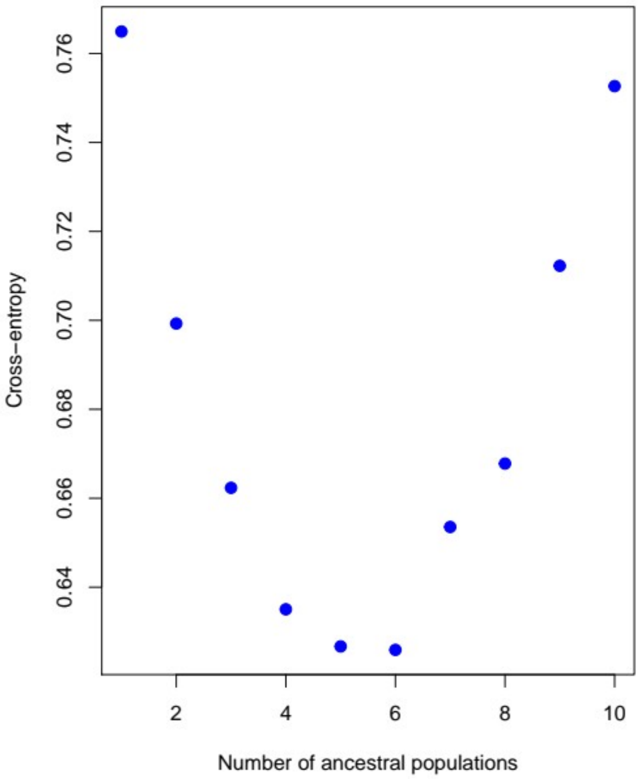

Fig. S1. LEA snmf Cross entropy estimates used to identify the number of latent factors for the latent factor mixed model gene-environment association analysis required to estimate the genetic offset.

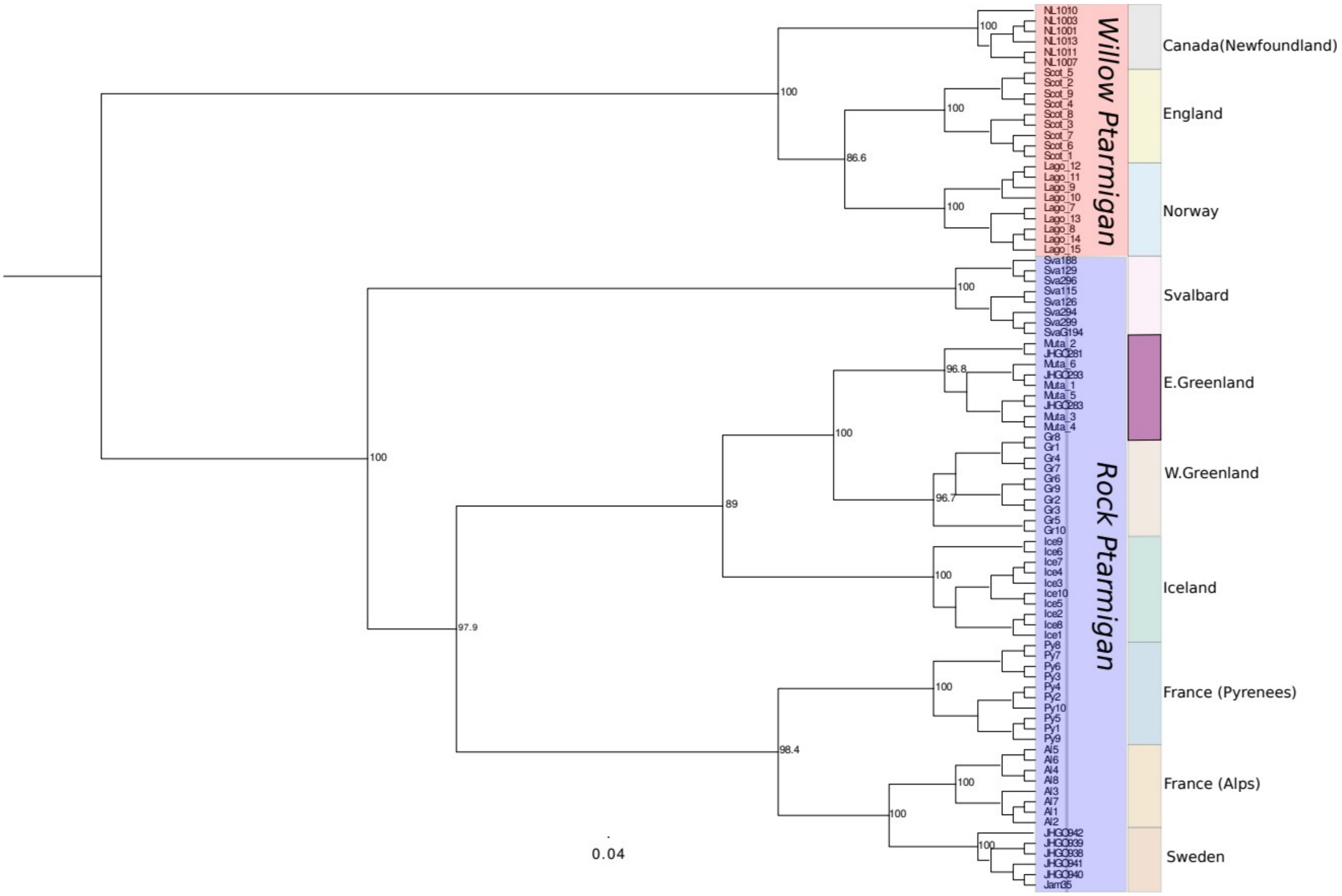

Fig. S2. Cladogram of maximum likelihood phylogenetic tree based on 3,205,910 variant sites, with support values from 1000 bootstrap replicates on the nodes.

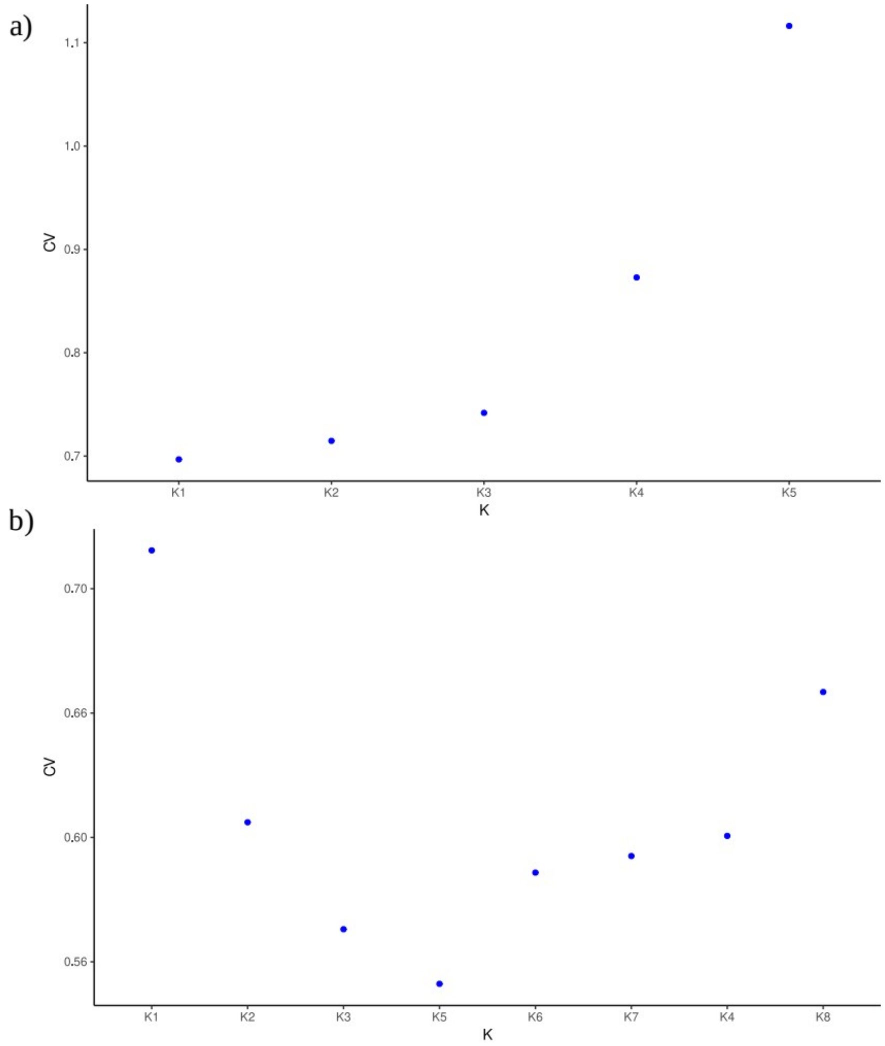

Fig. S3. Admixture cross validation scores to identify the optimal number of clusters (K) fitting the data. (a) willow ptarmigan, (b) rock ptarmigan.

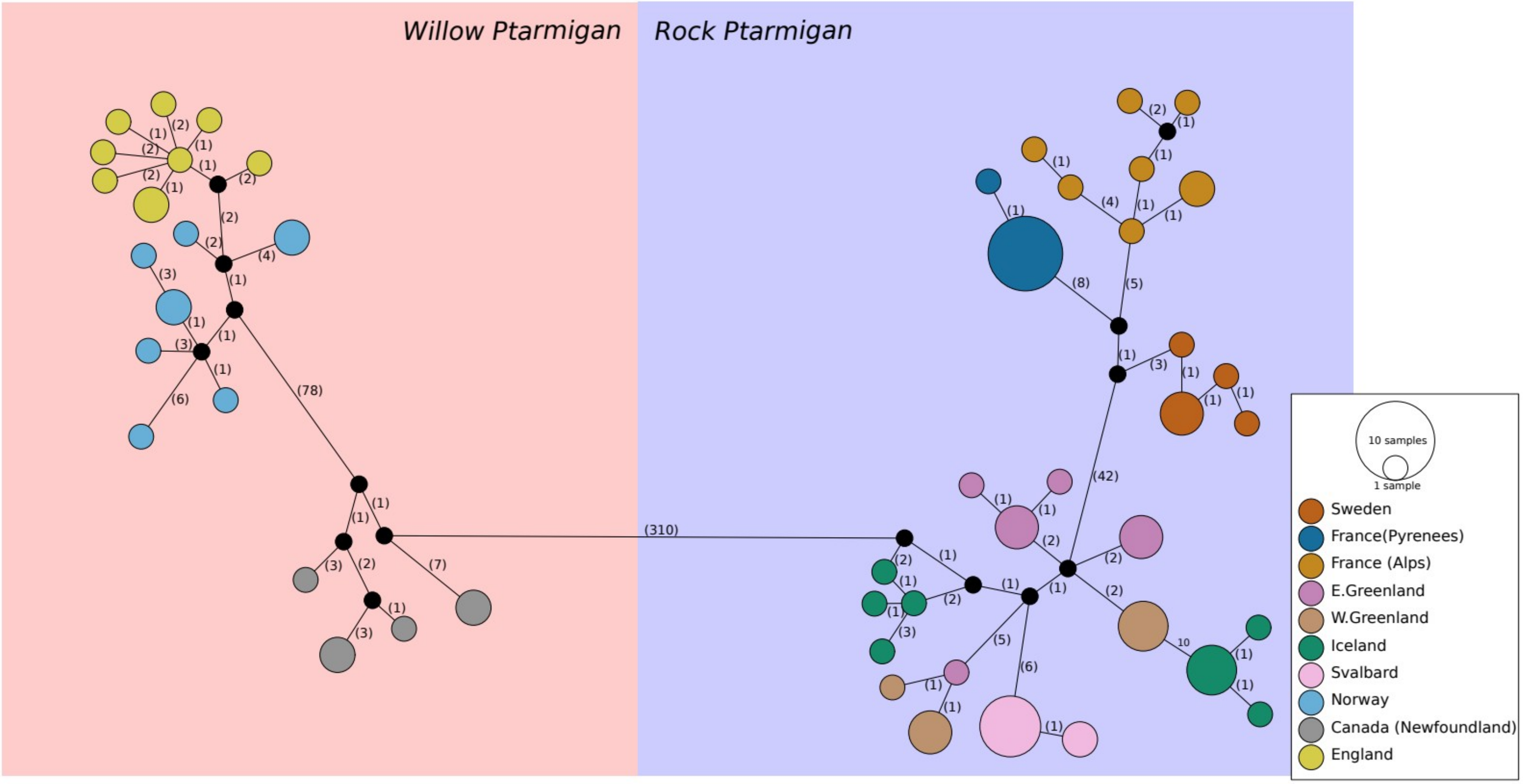

Fig. S4. Median joining mitochondria haplotype network. Numbers on connecting branches represent the number of mutational steps separating them and size of the circle represents the number of individuals with a specific haplotype.

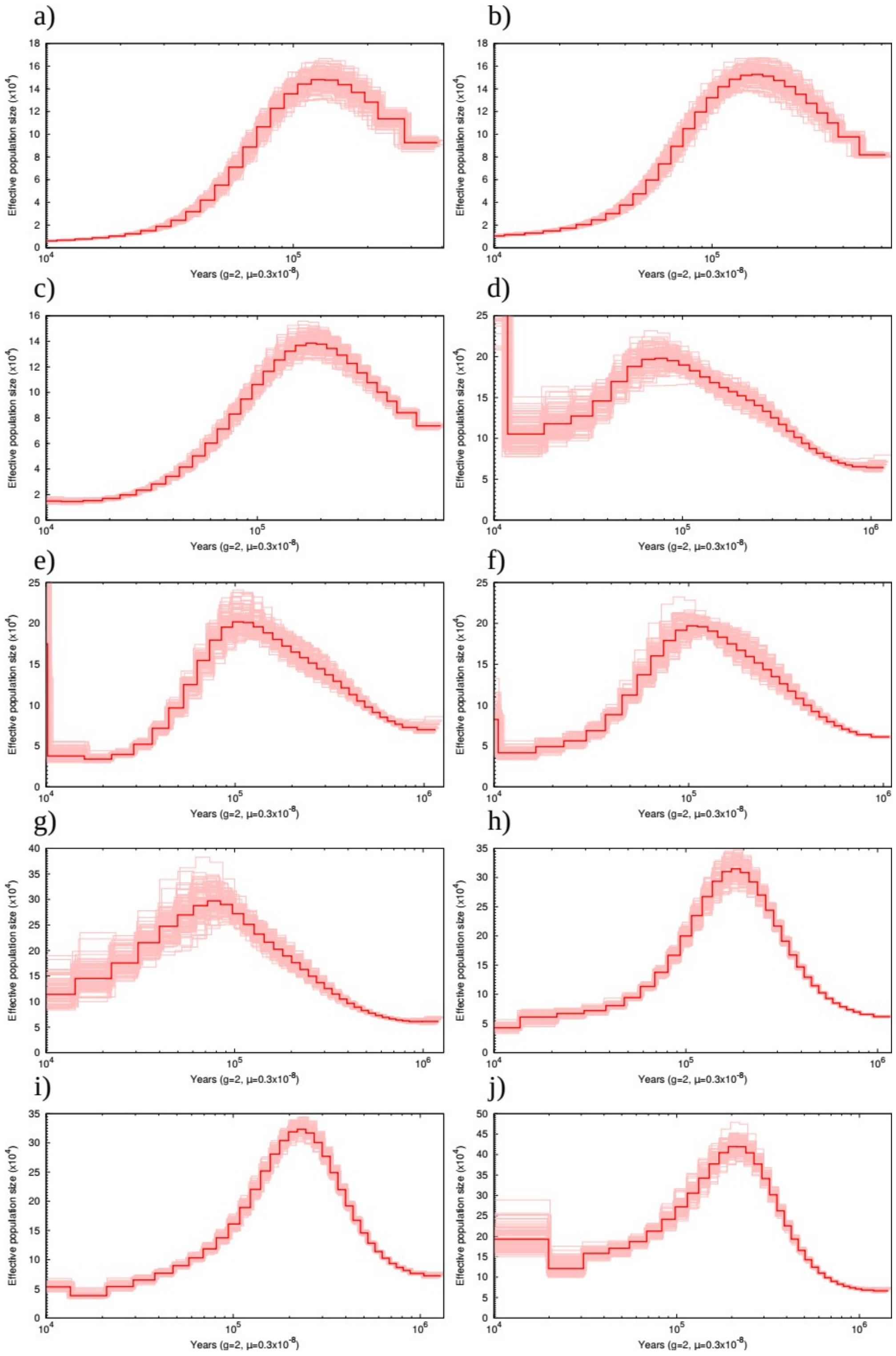

Fig. S5. PSMC based on 100 bootstrap replicates for (a) Svalbard, (b) France (Pyrenees), (c) Iceland, (d) Sweden, (e) Western Greenland, (f) Western Greenland, (g) France (Alps), (h) Canada (Newfoundland), (I) England, (j) Norway.

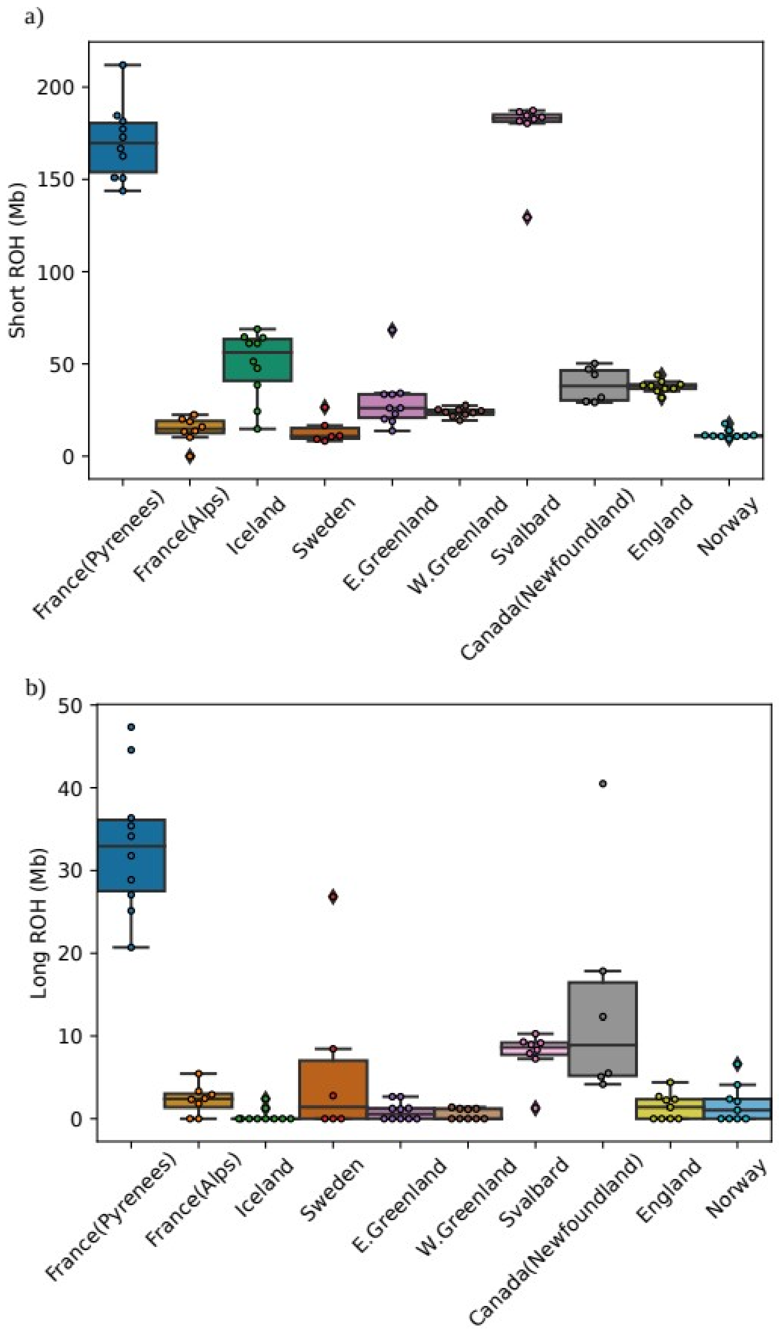

Fig. S6. The sum of runs of homozygosity (ROH) on a population level for (a) short ROH, (b) long ROH.

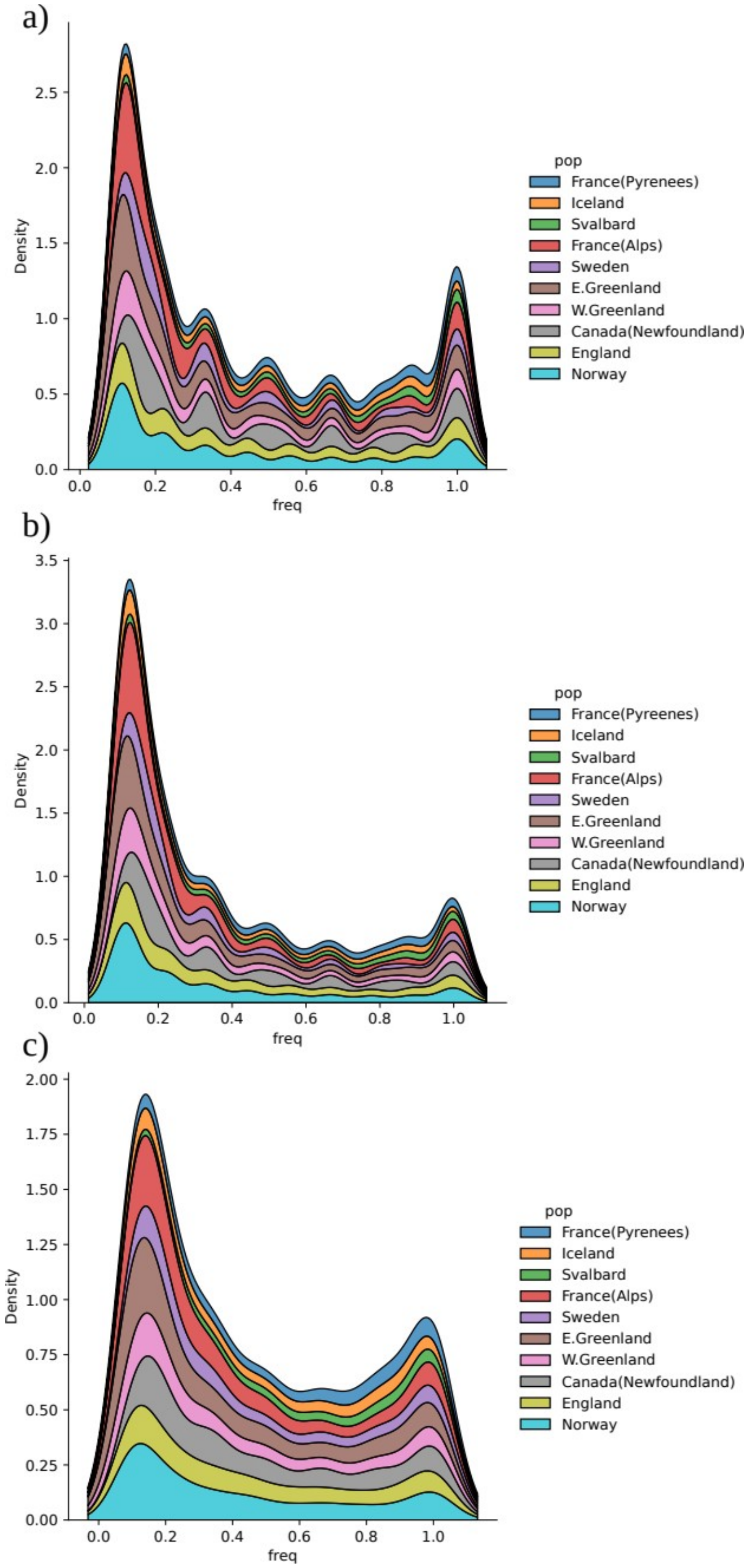

Fig. S7. Kernel density plot of allele frequency distribution of (a) missense mutations, (b) deleterious mutations, (c) loss-of-function mutations.

Table. S1. Harmonic mean effective population sizes. Nucleotide diversity (π) for autosomes and mitogenome and runs of homozygosity (ROH) based inbreeding coefficient ($F_{ROH}$) short and long ROH.

| Population | Ne(SNeP) | Ne(PSMC) | π(autosomes) | SD | π(mitogenome) | SD | FROH short | SD | FROH long | SD |
|---|---|---|---|---|---|---|---|---|---|---|
| France (Alps) | 157 | 223456 | 0.0032 | 0.0019 | 0.0218 | 0.0119 | 0.0150 | 0.0074 | 0.0024 | 0.0018 |
| E.Greenland | 208 | 130246 | 0.0026 | 0.0017 | 0.0492 | 0.0264 | 0.0310 | 0.0158 | 0.0009 | 0.0011 |
| WGreenland | 153 | 117962 | 0.0025 | 0.0016 | 0.0125 | 0.0069 | 0.0248 | 0.0024 | 0.0006 | 0.0007 |
| Iceland | 197 | 56716 | 0.0017 | 0.0014 | 0.0276 | 0.0146 | 0.0518 | 0.0192 | 0.0004 | 0.0008 |
| France (Pyrenees) | 154 | 56218 | 0.0015 | 0.0014 | 0.0000 | 0.0000 | 0.1772 | 0.0210 | 0.0345 | 0.0087 |
| Svalbard | 135 | 11925 | 0.0009 | 0.0011 | 0.0000 | 0.0000 | 0.1841 | 0.0202 | 0.0081 | 0.0029 |
| Sweden | 99 | 158652 | 0.0029 | 0.0024 | 0.0266 | 0.0154 | 0.0144 | 0.0072 | 0.0066 | 0.0110 |
| England | 190 | 141177 | 0.0036 | 0.0016 | 0.0087 | 0.0047 | 0.0393 | 0.0036 | 0.0015 | 0.0016 |
| Canada (Newfoundland) | 98 | 154904 | 0.0039 | 0.0017 | 0.0337 | 0.0195 | 0.0403 | 0.0099 | 0.0148 | 0.0145 |
| Norway | 151 | 265720 | 0.0043 | 0.0017 | 0.0240 | 0.0129 | 0.0125 | 0.0026 | 0.0019 | 0.0024 |

Table. S2. Allele frequency summary statistics for mutation load classes (missense, deleterious and loss-of-function).

**Missense mutations**

| pop | max | min | median | IQR | mean | SD |
|---|---|---|---|---|---|---|
| France (Pyrenees) | 1 | 0.1000 | 0.6000 | 0.6000 | 0.5814 | 0.3087 |
| Iceland | 1 | 0.1000 | 0.4000 | 0.7000 | 0.4759 | 0.3274 |
| Svalbard | 1 | 0.1250 | 0.6250 | 0.5000 | 0.6114 | 0.3124 |
| Fance (Alps) | 1 | 0.1250 | 0.2500 | 0.5000 | 0.3855 | 0.3111 |
| Sweden | 1 | 0.1667 | 0.3333 | 0.5000 | 0.4507 | 0.3118 |
| E.Greenland | 1 | 0.1000 | 0.3000 | 0.6000 | 0.3970 | 0.3216 |
| W.Greenland | 1 | 0.1111 | 0.3333 | 0.6667 | 0.4322 | 0.3296 |
| Canada (Newfoundland) | 1 | 0.1667 | 0.3333 | 0.6667 | 0.4853 | 0.3051 |
| England | 1 | 0.1111 | 0.3333 | 0.6667 | 0.4682 | 0.3210 |
| Norway | 1 | 0.1111 | 0.2222 | 0.5556 | 0.4081 | 0.3228 |

**Deleterious mutations**

| pop | max | min | median | IQR | mean | SD |
|---|---|---|---|---|---|---|
| France (Pyrenees) | 1 | 0.1000 | 0.5000 | 0.6000 | 0.5250 | 0.3149 |
| Iceland | 1 | 0.1000 | 0.2000 | 0.6000 | 0.3912 | 0.3191 |
| Svalbard | 1 | 0.1250 | 0.5000 | 0.6250 | 0.5593 | 0.3216 |
| Fance (Alps) | 1 | 0.1250 | 0.1250 | 0.2500 | 0.3145 | 0.2742 |
| Sweden | 1 | 0.1667 | 0.1667 | 0.3333 | 0.3900 | 0.2893 |
| E.Greenland | 1 | 0.1000 | 0.2000 | 0.4000 | 0.3285 | 0.2923 |
| W.Greenland | 1 | 0.1111 | 0.2222 | 0.4444 | 0.3601 | 0.3040 |
| Canada (Newfoundland) | 1 | 0.1667 | 0.3333 | 0.5000 | 0.4263 | 0.2901 |
| England | 1 | 0.1111 | 0.3333 | 0.5556 | 0.4050 | 0.3108 |
| Norway | 1 | 0.1111 | 0.2222 | 0.3333 | 0.3413 | 0.2935 |

**Loss-of-function mutations**

| pop | max | min | median | IQR | mean | SD |
|---|---|---|---|---|---|---|
| France (Pyrenees) | 1 | 0.1000 | 0.7000 | 0.6000 | 0.5943 | 0.3187 |
| Iceland | 1 | 0.1000 | 0.5000 | 0.6000 | 0.5056 | 0.3247 |
| Svalbard | 1 | 0.1250 | 0.6250 | 0.5000 | 0.6312 | 0.2918 |

| | | | | | | |
|---|---|---|---|---|---|---|
| Fance (Alps) | 1 | 0.1250 | 0.2500 | 0.5000 | 0.4060 | 0.3122 |
| Sweden | 1 | 0.1667 | 0.3333 | 0.6667 | 0.4896 | 0.3195 |
| E.Greenland | 1 | 0.1000 | 0.3000 | 0.6000 | 0.4022 | 0.3222 |
| W.Greenland | 1 | 0.1111 | 0.3333 | 0.6667 | 0.4429 | 0.3297 |
| Canada (Newfoundland) | 1 | 0.1667 | 0.3333 | 0.6667 | 0.4885 | 0.3042 |
| England | 1 | 0.1111 | 0.4444 | 0.5556 | 0.4768 | 0.3159 |
| Norway | 1 | 0.1111 | 0.3333 | 0.5556 | 0.4168 | 0.3183 |

Table. S3. Significantly enriched Gene ontology (GO) terms for each population, (a) missense mutations, (b) deleterious mutations, (c) loss-of-function mutations, (d) short runs of homozygosity ROH, (e) long ROH.

**a)**

**GO:Biological process**

| Pop | term_name | term_id | adjusted_p_value |
|---|---|---|---|
| Svalbard | double-strand break repair | GO:0006302 | 1.29E-06 |
| Svalbard | DNA repair | GO:0006281 | 3.02E-05 |
| Svalbard | cilium organization | GO:0044782 | 5.26E-04 |
| Svalbard | cilium assembly | GO:0060271 | 2.05E-03 |
| Svalbard | microtubule-based process | GO:0007017 | 2.28E-03 |
| Svalbard | DNA recombination | GO:0006310 | 2.64E-03 |
| Svalbard | meiotic chromosome segregation | GO:0045132 | 4.21E-03 |
| Svalbard | negative regulation of chromosome organization | GO:2001251 | 2.00E-02 |
| Svalbard | homologous chromosome segregation | GO:0045143 | 2.24E-02 |
| Svalbard | chromosome organization involved in meiotic cell cycle | GO:0070192 | 2.67E-02 |
| Svalbard | microtubule-based movement | GO:0007018 | 4.60E-02 |
| France (Pyrenees) | cilium organization | GO:0044782 | 3.03E-08 |
| France (Pyrenees) | double-strand break repair | GO:0006302 | 4.81E-08 |
| France (Pyrenees) | cilium assembly | GO:0060271 | 1.44E-07 |
| France (Pyrenees) | double-strand break repair via homologous recombination | GO:0000724 | 5.30E-04 |
| France (Pyrenees) | plasma membrane bounded cell projection assembly | GO:0120031 | 7.52E-04 |
| France (Pyrenees) | recombinational repair | GO:0000725 | 8.48E-04 |
| France (Pyrenees) | cell projection assembly | GO:0030031 | 1.85E-03 |
| France (Pyrenees) | DNA repair | GO:0006281 | 6.78E-03 |
| France (Pyrenees) | DNA recombination | GO:0006310 | 4.41E-02 |
| Iceland | double-strand break repair | GO:0006302 | 8.31E-10 |
| Iceland | cilium organization | GO:0044782 | 2.62E-09 |
| Iceland | cilium assembly | GO:0060271 | 1.11E-08 |
| Iceland | DNA repair | GO:0006281 | 1.47E-06 |
| Iceland | organelle assembly | GO:0070925 | 7.84E-05 |
| Iceland | plasma membrane bounded cell projection assembly | GO:0120031 | 9.32E-05 |
| Iceland | microtubule-based process | GO:0007017 | 1.79E-04 |
| Iceland | cell projection assembly | GO:0030031 | 1.97E-04 |
| Iceland | non-recombinational repair | GO:0000726 | 2.86E-04 |

| | | | |
|---|---|---|---|
| Iceland | cellular response to DNA damage stimulus | GO:0006974 | 7.75E-04 |
| Iceland | centriole assembly | GO:0098534 | 1.28E-03 |
| Iceland | centriole replication | GO:0007099 | 3.58E-03 |
| Iceland | DNA recombination | GO:0006310 | 3.83E-03 |
| Iceland | meiotic chromosome segregation | GO:0045132 | 5.23E-03 |
| Iceland | double-strand break repair via nonhomologous end joining | GO:0006303 | 5.92E-03 |
| Iceland | chromosome segregation | GO:0007059 | 6.22E-03 |
| Iceland | chromosome organization involved in meiotic cell cycle | GO:0070192 | 8.30E-03 |
| Iceland | microtubule cytoskeleton organization | GO:0000226 | 1.08E-02 |
| Iceland | microtubule-based movement | GO:0007018 | 1.09E-02 |
| Iceland | centrosome duplication | GO:0051298 | 1.14E-02 |
| Iceland | nuclear chromosome segregation | GO:0098813 | 1.50E-02 |
| Iceland | microtubule organizing center organization | GO:0031023 | 1.71E-02 |
| Iceland | centrosome cycle | GO:0007098 | 2.18E-02 |
| Iceland | homologous chromosome segregation | GO:0045143 | 2.64E-02 |
| Iceland | meiotic cell cycle process | GO:1903046 | 4.17E-02 |
| E.Greenland | DNA repair | GO:0006281 | 2.13E-07 |
| E.Greenland | double-strand break repair | GO:0006302 | 3.42E-07 |
| E.Greenland | cilium organization | GO:0044782 | 2.04E-06 |
| E.Greenland | cilium assembly | GO:0060271 | 7.01E-06 |
| E.Greenland | cellular response to DNA damage stimulus | GO:0006974 | 8.41E-06 |
| E.Greenland | microtubule-based process | GO:0007017 | 2.52E-03 |
| E.Greenland | axoneme assembly | GO:0035082 | 2.80E-03 |
| E.Greenland | DNA replication | GO:0006260 | 7.16E-03 |
| E.Greenland | DNA recombination | GO:0006310 | 8.69E-03 |
| E.Greenland | non-recombinational repair | GO:0000726 | 9.91E-03 |
| E.Greenland | cellular amino acid metabolic process | GO:0006520 | 1.87E-02 |
| E.Greenland | regulation of DNA replication | GO:0006275 | 2.29E-02 |
| E.Greenland | microtubule-based movement | GO:0007018 | 3.78E-02 |
| W.Greenland | cilium organization | GO:0044782 | 1.80E-09 |
| W.Greenland | double-strand break repair | GO:0006302 | 7.23E-09 |
| W.Greenland | cilium assembly | GO:0060271 | 2.35E-08 |
| W.Greenland | DNA repair | GO:0006281 | 2.55E-06 |
| W.Greenland | microtubule-based process | GO:0007017 | 1.11E-05 |
| W.Greenland | cell cycle process | GO:0022402 | 1.76E-05 |
| W.Greenland | microtubule cytoskeleton organization | GO:0000226 | 2.15E-05 |

| | | | |
|---|---|---|---|
| W.Greenland | chromosome segregation | GO:0007059 | 2.41E-05 |
| W.Greenland | chromosome organization involved in meiotic cell cycle | GO:0070192 | 8.33E-05 |
| W.Greenland | cellular response to DNA damage stimulus | GO:0006974 | 1.38E-04 |
| W.Greenland | plasma membrane bounded cell projection assembly | GO:0120031 | 1.76E-04 |
| W.Greenland | DNA replication | GO:0006260 | 2.13E-04 |
| W.Greenland | cell projection assembly | GO:0030031 | 2.48E-04 |
| W.Greenland | meiotic chromosome segregation | GO:0045132 | 2.53E-04 |
| W.Greenland | double-strand break repair via homologous recombination | GO:0000724 | 9.86E-04 |
| W.Greenland | cell cycle | GO:0007049 | 1.16E-03 |
| W.Greenland | recombinational repair | GO:0000725 | 1.55E-03 |
| W.Greenland | nuclear chromosome segregation | GO:0098813 | 2.20E-03 |
| W.Greenland | non-motile cilium assembly | GO:1905515 | 2.25E-03 |
| W.Greenland | axoneme assembly | GO:0035082 | 4.15E-03 |
| W.Greenland | nuclear division | GO:0000280 | 5.44E-03 |
| W.Greenland | sister chromatid segregation | GO:0000819 | 5.82E-03 |
| W.Greenland | DNA recombination | GO:0006310 | 6.24E-03 |
| W.Greenland | organelle assembly | GO:0070925 | 6.56E-03 |
| W.Greenland | organelle fission | GO:0048285 | 1.17E-02 |
| W.Greenland | homologous chromosome segregation | GO:0045143 | 1.20E-02 |
| W.Greenland | photoreceptor cell maintenance | GO:0045494 | 1.51E-02 |
| France(Alps) | double-strand break repair | GO:0006302 | 1.09E-07 |
| France(Alps) | DNA repair | GO:0006281 | 1.98E-07 |
| France(Alps) | cilium organization | GO:0044782 | 1.13E-06 |
| France(Alps) | DNA recombination | GO:0006310 | 5.84E-06 |
| France(Alps) | cellular response to DNA damage stimulus | GO:0006974 | 8.88E-06 |
| France(Alps) | microtubule-based process | GO:0007017 | 2.51E-05 |
| France(Alps) | cilium assembly | GO:0060271 | 3.09E-05 |
| France(Alps) | meiotic chromosome segregation | GO:0045132 | 1.24E-04 |
| France(Alps) | chromosome organization involved in meiotic cell cycle | GO:0070192 | 2.81E-04 |
| France(Alps) | microtubule-based movement | GO:0007018 | 4.60E-04 |
| France(Alps) | centrosome cycle | GO:0007098 | 8.88E-04 |
| France(Alps) | microtubule cytoskeleton organization | GO:0000226 | 9.77E-04 |
| France(Alps) | chromosome segregation | GO:0007059 | 1.19E-03 |
| France(Alps) | microtubule organizing center organization | GO:0031023 | 2.02E-03 |
| France(Alps) | centrosome duplication | GO:0051298 | 3.76E-03 |
| France(Alps) | meiotic nuclear division | GO:0140013 | 1.37E-02 |

| | | | |
|---|---|---|---|
| France(Alps) | meiotic cell cycle process | GO:1903046 | 1.60E-02 |
| France(Alps) | glycerolipid metabolic process | GO:0046486 | 1.72E-02 |
| France(Alps) | homologous chromosome segregation | GO:0045143 | 1.81E-02 |
| France(Alps) | DNA double-strand break processing | GO:0000729 | 2.02E-02 |
| France(Alps) | cell cycle process | GO:0022402 | 2.47E-02 |
| France(Alps) | meiotic cell cycle | GO:0051321 | 2.85E-02 |
| France(Alps) | phospholipid metabolic process | GO:0006644 | 3.15E-02 |
| France(Alps) | cell projection assembly | GO:0030031 | 3.50E-02 |
| France(Alps) | meiosis I cell cycle process | GO:0061982 | 3.63E-02 |
| France(Alps) | meiosis I | GO:0007127 | 3.63E-02 |
| France(Alps) | plasma membrane bounded cell projection assembly | GO:0120031 | 3.70E-02 |
| France(Alps) | axoneme assembly | GO:0035082 | 4.32E-02 |
| France(Alps) | cilium movement | GO:0003341 | 4.50E-02 |
| Sweden | double-strand break repair | GO:0006302 | 4.47E-10 |
| Sweden | DNA repair | GO:0006281 | 1.05E-06 |
| Sweden | DNA recombination | GO:0006310 | 5.19E-05 |
| Sweden | non-recombinational repair | GO:0000726 | 5.55E-05 |
| Sweden | cellular response to DNA damage stimulus | GO:0006974 | 6.57E-05 |
| Sweden | microtubule-based process | GO:0007017 | 1.22E-04 |
| Sweden | cilium assembly | GO:0060271 | 1.37E-04 |
| Sweden | cilium organization | GO:0044782 | 1.95E-04 |
| Sweden | double-strand break repair via nonhomologous end joining | GO:0006303 | 1.04E-03 |
| Sweden | meiotic chromosome segregation | GO:0045132 | 2.33E-03 |
| Sweden | microtubule-based movement | GO:0007018 | 6.28E-03 |
| Sweden | organelle assembly | GO:0070925 | 9.20E-03 |
| Sweden | cell cycle process | GO:0022402 | 1.37E-02 |
| Sweden | negative regulation of DNA metabolic process | GO:0051053 | 1.56E-02 |
| Sweden | double-strand break repair via homologous recombination | GO:0000724 | 2.99E-02 |
| Sweden | microtubule cytoskeleton organization | GO:0000226 | 3.18E-02 |
| Sweden | cilium movement | GO:0003341 | 3.26E-02 |
| Sweden | chromosome organization involved in meiotic cell cycle | GO:0070192 | 3.65E-02 |
| Sweden | recombinational repair | GO:0000725 | 4.21E-02 |
| Sweden | DNA-dependent DNA replication maintenance of fidelity | GO:0045005 | 4.49E-02 |
| England | double-strand break repair | GO:0006302 | 2.02E-10 |
| England | microtubule-based process | GO:0007017 | 1.33E-09 |
| England | cilium organization | GO:0044782 | 4.50E-09 |

| | | | |
|---|---|---|---|
| England | cilium assembly | GO:0060271 | 1.65E-08 |
| England | DNA repair | GO:0006281 | 4.90E-08 |
| England | microtubule cytoskeleton organization | GO:0000226 | 4.87E-07 |
| England | cellular response to DNA damage stimulus | GO:0006974 | 7.18E-07 |
| England | DNA recombination | GO:0006310 | 7.25E-05 |
| England | double-strand break repair via homologous recombination | GO:0000724 | 4.56E-04 |
| England | microtubule-based movement | GO:0007018 | 6.47E-04 |
| England | recombinational repair | GO:0000725 | 7.46E-04 |
| England | DNA replication | GO:0006260 | 1.04E-03 |
| England | meiotic chromosome segregation | GO:0045132 | 1.87E-03 |
| England | chromosome organization involved in meiotic cell cycle | GO:0070192 | 2.41E-03 |
| England | cell cycle process | GO:0022402 | 2.69E-03 |
| England | chromosome segregation | GO:0007059 | 2.84E-03 |
| England | plasma membrane bounded cell projection assembly | GO:0120031 | 3.66E-03 |
| England | axoneme assembly | GO:0035082 | 5.02E-03 |
| England | cell projection assembly | GO:0030031 | 5.29E-03 |
| England | nuclear chromosome segregation | GO:0098813 | 7.19E-03 |
| England | protein import into peroxisome matrix | GO:0016558 | 7.22E-03 |
| England | replication fork processing | GO:0031297 | 8.77E-03 |
| England | DNA-dependent DNA replication | GO:0006261 | 9.35E-03 |
| England | organelle assembly | GO:0070925 | 9.79E-03 |
| England | DNA-dependent DNA replication maintenance of fidelity | GO:0045005 | 1.04E-02 |
| England | meiotic nuclear division | GO:0140013 | 1.33E-02 |
| England | microtubule bundle formation | GO:0001578 | 3.09E-02 |
| England | cilium movement | GO:0003341 | 3.46E-02 |
| England | protein localization to peroxisome | GO:0072662 | 3.94E-02 |
| England | establishment of protein localization to peroxisome | GO:0072663 | 3.94E-02 |
| England | protein targeting to peroxisome | GO:0006625 | 3.94E-02 |
| England | regulation of DNA replication | GO:0006275 | 4.39E-02 |
| England | homologous chromosome segregation | GO:0045143 | 4.73E-02 |
| England | mitotic cell cycle checkpoint | GO:0007093 | 4.93E-02 |
| Canada (Newfoundland) | double-strand break repair | GO:0006302 | 1.54E-07 |
| Canada (Newfoundland) | cilium organization | GO:0044782 | 1.21E-06 |
| Canada (Newfoundland) | cilium assembly | GO:0060271 | 1.04E-05 |
| Canada (Newfoundland) | microtubule-based process | GO:0007017 | 1.34E-05 |
| Canada (Newfoundland) | DNA replication | GO:0006260 | 5.73E-05 |

| | | | |
|---|---|---|---|
| Canada (Newfoundland) | DNA repair | GO:0006281 | 8.05E-05 |
| Canada (Newfoundland) | microtubule cytoskeleton organization | GO:0000226 | 1.00E-04 |
| Canada (Newfoundland) | cellular response to DNA damage stimulus | GO:0006974 | 9.35E-04 |
| Canada (Newfoundland) | microtubule bundle formation | GO:0001578 | 3.54E-03 |
| Canada (Newfoundland) | axoneme assembly | GO:0035082 | 5.52E-03 |
| Canada (Newfoundland) | ncRNA metabolic process | GO:0034660 | 5.72E-03 |
| Canada (Newfoundland) | replication fork processing | GO:0031297 | 6.31E-03 |
| Canada (Newfoundland) | anatomical structure homeostasis | GO:0060249 | 6.43E-03 |
| Canada (Newfoundland) | microtubule-based movement | GO:0007018 | 7.44E-03 |
| Canada (Newfoundland) | DNA recombination | GO:0006310 | 8.48E-03 |
| Canada (Newfoundland) | DNA-dependent DNA replication maintenance of fidelity | GO:0045005 | 1.29E-02 |
| Canada (Newfoundland) | double-strand break repair via homologous recombination | GO:0000724 | 1.35E-02 |
| Canada (Newfoundland) | plasma membrane bounded cell projection assembly | GO:0120031 | 1.67E-02 |
| Canada (Newfoundland) | cellular amino acid metabolic process | GO:0006520 | 1.90E-02 |
| Canada (Newfoundland) | DNA-dependent DNA replication | GO:0006261 | 2.15E-02 |
| Canada (Newfoundland) | recombinational repair | GO:0000725 | 2.16E-02 |
| Canada (Newfoundland) | tRNA metabolic process | GO:0006399 | 2.71E-02 |
| Canada (Newfoundland) | regulation of DNA replication | GO:0006275 | 2.81E-02 |
| Canada (Newfoundland) | cell projection assembly | GO:0030031 | 4.66E-02 |
| Norway | double-strand break repair | GO:0006302 | 7.94E-10 |
| Norway | cilium organization | GO:0044782 | 1.48E-09 |
| Norway | cilium assembly | GO:0060271 | 1.73E-08 |
| Norway | DNA repair | GO:0006281 | 1.17E-07 |
| Norway | DNA recombination | GO:0006310 | 6.75E-06 |
| Norway | cellular response to DNA damage stimulus | GO:0006974 | 6.67E-05 |
| Norway | microtubule-based process | GO:0007017 | 1.74E-04 |
| Norway | meiotic chromosome segregation | GO:0045132 | 2.12E-04 |
| Norway | double-strand break repair via homologous recombination | GO:0000724 | 2.22E-04 |
| Norway | recombinational repair | GO:0000725 | 3.96E-04 |
| Norway | microtubule cytoskeleton organization | GO:0000226 | 2.04E-03 |
| Norway | non-recombinational repair | GO:0000726 | 2.49E-03 |
| Norway | plasma membrane bounded cell projection assembly | GO:0120031 | 2.64E-03 |
| Norway | cell projection assembly | GO:0030031 | 2.79E-03 |
| Norway | replication fork processing | GO:0031297 | 4.02E-03 |
| Norway | non-motile cilium assembly | GO:1905515 | 4.89E-03 |
| Norway | DNA replication | GO:0006260 | 5.34E-03 |

| | | | |
|---|---|---|---|
| Norway | chromosome organization involved in meiotic cell cycle | GO:0070192 | 6.67E-03 |
| Norway | DNA-dependent DNA replication maintenance of fidelity | GO:0045005 | 8.01E-03 |
| Norway | axoneme assembly | GO:0035082 | 1.26E-02 |
| Norway | microtubule bundle formation | GO:0001578 | 1.47E-02 |
| Norway | microtubule-based movement | GO:0007018 | 1.74E-02 |
| Norway | organelle assembly | GO:0070925 | 2.86E-02 |
| Norway | DNA double-strand break processing | GO:0000729 | 4.06E-02 |

**GO:Molecular function**

| Pop | term_name | term_id | adjusted_p_value |
|---|---|---|---|
| Svalbard | ATPase activity | GO:0016887 | 3.00E-05 |
| Svalbard | motor activity | GO:0003774 | 5.91E-04 |
| Svalbard | guanyl-nucleotide exchange factor activity | GO:0005085 | 3.62E-03 |
| France (Pyrenees) | ATPase activity | GO:0016887 | 2.23E-05 |
| France (Pyrenees) | adenyl ribonucleotide binding | GO:0032559 | 7.18E-05 |
| France (Pyrenees) | ATP binding | GO:0005524 | 1.11E-04 |
| France (Pyrenees) | adenyl nucleotide binding | GO:0030554 | 1.15E-04 |
| France (Pyrenees) | catalytic activity | GO:0003824 | 1.06E-03 |
| France (Pyrenees) | motor activity | GO:0003774 | 1.46E-02 |
| France (Pyrenees) | hydrolase activity, acting on glycosyl bonds | GO:0016798 | 2.65E-02 |
| France (Pyrenees) | anion binding | GO:0043168 | 2.80E-02 |
| France (Pyrenees) | ion binding | GO:0043167 | 4.31E-02 |
| Iceland | ATPase activity | GO:0016887 | 1.91E-05 |
| Iceland | motor activity | GO:0003774 | 1.79E-03 |
| Iceland | protein binding | GO:0005515 | 4.01E-03 |
| Iceland | ATP binding | GO:0005524 | 2.50E-02 |
| E.Greenland | catalytic activity | GO:0003824 | 2.33E-07 |
| E.Greenland | oxidoreductase activity, acting on paired donors, with incorporation or reduction of molecular oxygen | GO:0016705 | 4.75E-05 |
| E.Greenland | hydrolase activity | GO:0016787 | 9.41E-05 |
| E.Greenland | ATPase activity | GO:0016887 | 5.03E-04 |
| E.Greenland | metalloendopeptidase activity | GO:0004222 | 1.33E-03 |
| E.Greenland | immune receptor activity | GO:0140375 | 3.84E-03 |
| E.Greenland | metallopeptidase activity | GO:0008237 | 4.20E-03 |
| E.Greenland | transferase activity, transferring one-carbon groups | GO:0016741 | 5.31E-03 |
| E.Greenland | methyltransferase activity | GO:0008168 | 8.46E-03 |
| E.Greenland | hydrolase activity, acting on glycosyl bonds | GO:0016798 | 1.41E-02 |

| | | | |
|---|---|---|---|
| E.Greenland | adenyl nucleotide binding | GO:0030554 | 1.59E-02 |
| E.Greenland | cytokine receptor activity | GO:0004896 | 1.67E-02 |
| E.Greenland | tetrapyrrole binding | GO:0046906 | 1.94E-02 |
| E.Greenland | guanyl-nucleotide exchange factor activity | GO:0005085 | 1.97E-02 |
| E.Greenland | adenyl ribonucleotide binding | GO:0032559 | 2.07E-02 |
| E.Greenland | hydrolase activity, hydrolyzing O-glycosyl compounds | GO:0004553 | 2.07E-02 |
| E.Greenland | iron ion binding | GO:0005506 | 2.80E-02 |
| E.Greenland | ligase activity | GO:0016874 | 3.08E-02 |
| E.Greenland | ATP binding | GO:0005524 | 3.80E-02 |
| E.Greenland | modification-dependent protein binding | GO:0140030 | 4.54E-02 |
| E.Greenland | microtubule motor activity | GO:0003777 | 4.93E-02 |
| W.Greenland | ATPase activity | GO:0016887 | 1.32E-07 |
| W.Greenland | adenyl nucleotide binding | GO:0030554 | 3.15E-04 |
| W.Greenland | ATP binding | GO:0005524 | 3.20E-04 |
| W.Greenland | adenyl ribonucleotide binding | GO:0032559 | 3.95E-04 |
| W.Greenland | catalytic activity | GO:0003824 | 8.42E-04 |
| W.Greenland | motor activity | GO:0003774 | 8.73E-04 |
| W.Greenland | metalloendopeptidase activity | GO:0004222 | 8.86E-04 |
| W.Greenland | hydrolase activity | GO:0016787 | 1.22E-03 |
| W.Greenland | guanyl-nucleotide exchange factor activity | GO:0005085 | 4.23E-03 |
| W.Greenland | catalytic activity, acting on DNA | GO:0140097 | 5.49E-03 |
| W.Greenland | microtubule motor activity | GO:0003777 | 1.41E-02 |
| W.Greenland | protein-glutamine gamma-glutamyltransferase activity | GO:0003810 | 2.26E-02 |
| W.Greenland | cytokine receptor activity | GO:0004896 | 2.41E-02 |
| W.Greenland | deoxyribonuclease activity | GO:0004536 | 2.46E-02 |
| W.Greenland | immune receptor activity | GO:0140375 | 2.55E-02 |
| W.Greenland | metallopeptidase activity | GO:0008237 | 2.58E-02 |
| W.Greenland | tRNA methyltransferase activity | GO:0008175 | 3.67E-02 |
| W.Greenland | methyltransferase activity | GO:0008168 | 4.94E-02 |
| France(Alps) | catalytic activity | GO:0003824 | 4.31E-09 |
| France(Alps) | ATPase activity | GO:0016887 | 8.80E-08 |
| France(Alps) | ATP binding | GO:0005524 | 1.79E-06 |
| France(Alps) | adenyl nucleotide binding | GO:0030554 | 2.12E-06 |
| France(Alps) | adenyl ribonucleotide binding | GO:0032559 | 2.31E-06 |
| France(Alps) | motor activity | GO:0003774 | 7.77E-06 |
| France(Alps) | hydrolase activity | GO:0016787 | 9.73E-05 |

| | | | |
|---|---|---|---|
| France(Alps) | metalloendopeptidase activity | GO:0004222 | 1.46E-03 |
| France(Alps) | metallopeptidase activity | GO:0008237 | 2.25E-03 |
| France(Alps) | DNA helicase activity | GO:0003678 | 4.01E-03 |
| France(Alps) | guanyl-nucleotide exchange factor activity | GO:0005085 | 4.03E-03 |
| France(Alps) | oxidoreductase activity, acting on paired donors, with incorporation or reduction of molecular oxygen | GO:0016705 | 5.19E-03 |
| France(Alps) | catalytic activity, acting on DNA | GO:0140097 | 8.53E-03 |
| France(Alps) | ion binding | GO:0043167 | 1.03E-02 |
| France(Alps) | iron ion binding | GO:0005506 | 1.21E-02 |
| France(Alps) | hydrolase activity, acting on glycosyl bonds | GO:0016798 | 2.31E-02 |
| France(Alps) | calcium ion binding | GO:0005509 | 2.50E-02 |
| France(Alps) | hydrolase activity, hydrolyzing O-glycosyl compounds | GO:0004553 | 3.63E-02 |
| France(Alps) | microtubule motor activity | GO:0003777 | 4.78E-02 |
| Sweden | ATPase activity | GO:0016887 | 2.77E-05 |
| Sweden | adenyl nucleotide binding | GO:0030554 | 1.43E-03 |
| Sweden | ATP binding | GO:0005524 | 1.71E-03 |
| Sweden | adenyl ribonucleotide binding | GO:0032559 | 1.94E-03 |
| Sweden | motor activity | GO:0003774 | 1.52E-02 |
| Sweden | microtubule motor activity | GO:0003777 | 1.75E-02 |
| Sweden | extracellular matrix structural constituent | GO:0005201 | 1.94E-02 |
| Sweden | carboxylic ester hydrolase activity | GO:0052689 | 4.03E-02 |
| England | ATPase activity | GO:0016887 | 8.30E-08 |
| England | catalytic activity | GO:0003824 | 4.22E-06 |
| England | hydrolase activity | GO:0016787 | 4.58E-04 |
| England | catalytic activity, acting on DNA | GO:0140097 | 2.11E-03 |
| England | guanyl-nucleotide exchange factor activity | GO:0005085 | 5.90E-03 |
| England | ATP-dependent microtubule motor activity | GO:1990939 | 7.28E-03 |
| England | ATP binding | GO:0005524 | 7.34E-03 |
| England | protein binding | GO:0005515 | 9.53E-03 |
| England | DNA helicase activity | GO:0003678 | 1.07E-02 |
| England | peptidase regulator activity | GO:0061134 | 1.09E-02 |
| England | adenyl ribonucleotide binding | GO:0032559 | 1.43E-02 |
| England | microtubule binding | GO:0008017 | 1.51E-02 |
| England | microtubule motor activity | GO:0003777 | 1.59E-02 |
| England | adenyl nucleotide binding | GO:0030554 | 1.63E-02 |
| England | deoxyribonuclease activity | GO:0004536 | 2.59E-02 |
| England | motor activity | GO:0003774 | 3.47E-02 |

| | | | |
|---|---|---|---|
| England | cysteine-type peptidase activity | GO:0008234 | 4.09E-02 |
| England | hydrolase activity, acting on glycosyl bonds | GO:0016798 | 4.72E-02 |
| Canada (Newfoundland) | ATP binding | GO:0005524 | 3.12E-05 |
| Canada (Newfoundland) | catalytic activity | GO:0003824 | 3.32E-05 |
| Canada (Newfoundland) | adenyl nucleotide binding | GO:0030554 | 3.89E-05 |
| Canada (Newfoundland) | adenyl ribonucleotide binding | GO:0032559 | 3.91E-05 |
| Canada (Newfoundland) | ATPase activity | GO:0016887 | 5.85E-05 |
| Canada (Newfoundland) | guanyl-nucleotide exchange factor activity | GO:0005085 | 9.74E-05 |
| Canada (Newfoundland) | RNA methyltransferase activity | GO:0008173 | 2.05E-03 |
| Canada (Newfoundland) | protein binding | GO:0005515 | 2.25E-03 |
| Canada (Newfoundland) | motor activity | GO:0003774 | 2.40E-03 |
| Canada (Newfoundland) | tubulin binding | GO:0015631 | 2.65E-03 |
| Canada (Newfoundland) | methyltransferase activity | GO:0008168 | 3.02E-03 |
| Canada (Newfoundland) | transferase activity, transferring one-carbon groups | GO:0016741 | 3.65E-03 |
| Canada (Newfoundland) | microtubule binding | GO:0008017 | 3.79E-03 |
| Canada (Newfoundland) | S-adenosylmethionine-dependent methyltransferase activity | GO:0008757 | 6.16E-03 |
| Canada (Newfoundland) | deoxyribonuclease activity | GO:0004536 | 6.64E-03 |
| Canada (Newfoundland) | anion binding | GO:0043168 | 1.23E-02 |
| Canada (Newfoundland) | active transmembrane transporter activity | GO:0022804 | 1.44E-02 |
| Canada (Newfoundland) | endodeoxyribonuclease activity | GO:0004520 | 1.67E-02 |
| Canada (Newfoundland) | catalytic activity, acting on DNA | GO:0140097 | 2.26E-02 |
| Canada (Newfoundland) | small molecule binding | GO:0036094 | 3.07E-02 |
| Canada (Newfoundland) | catalytic activity, acting on a tRNA | GO:0140101 | 3.85E-02 |
| Canada (Newfoundland) | carbohydrate derivative binding | GO:0097367 | 4.02E-02 |
| Canada (Newfoundland) | protein-lysine N-methyltransferase activity | GO:0016279 | 4.32E-02 |
| Canada (Newfoundland) | lysine N-methyltransferase activity | GO:0016278 | 4.32E-02 |
| Canada (Newfoundland) | cytoskeletal protein binding | GO:0008092 | 4.35E-02 |
| Canada (Newfoundland) | nucleoside phosphate binding | GO:1901265 | 4.38E-02 |
| Canada (Newfoundland) | nucleotide binding | GO:0000166 | 4.38E-02 |
| Norway | catalytic activity | GO:0003824 | 1.10E-10 |
| Norway | ATPase activity | GO:0016887 | 6.55E-09 |
| Norway | guanyl-nucleotide exchange factor activity | GO:0005085 | 9.37E-06 |
| Norway | hydrolase activity | GO:0016787 | 3.35E-05 |
| Norway | adenyl ribonucleotide binding | GO:0032559 | 7.43E-05 |
| Norway | adenyl nucleotide binding | GO:0030554 | 1.00E-04 |
| Norway | ATP binding | GO:0005524 | 1.33E-04 |

| | | | |
|---|---|---|---|
| Norway | iron ion binding | GO:0005506 | 7.58E-04 |
| Norway | deoxyribonuclease activity | GO:0004536 | 3.75E-03 |
| Norway | motor activity | GO:0003774 | 4.23E-03 |
| Norway | oxidoreductase activity, acting on paired donors, with incorporation or reduction of molecular oxygen | GO:0016705 | 5.00E-03 |
| Norway | transferase activity, transferring glycosyl groups | GO:0016757 | 5.71E-03 |
| Norway | catalytic activity, acting on DNA | GO:0140097 | 6.79E-03 |
| Norway | metallopeptidase activity | GO:0008237 | 1.42E-02 |
| Norway | transferase activity, transferring one-carbon groups | GO:0016741 | 2.86E-02 |
| Norway | cytokine receptor activity | GO:0004896 | 3.39E-02 |
| Norway | ATP-dependent microtubule motor activity | GO:1990939 | 4.98E-02 |

**GO:Cellular component**

| Pop | term_name | term_id | adjusted_p_value |
|---|---|---|---|
| Svalbard | microtubule cytoskeleton | GO:0015630 | 4.31E-04 |
| Svalbard | ciliary plasm | GO:0097014 | 6.47E-04 |
| Svalbard | axoneme | GO:0005930 | 1.37E-03 |
| Svalbard | centrosome | GO:0005813 | 1.90E-03 |
| Svalbard | microtubule organizing center | GO:0005815 | 2.05E-03 |
| Svalbard | cilium | GO:0005929 | 7.19E-03 |
| Svalbard | synaptonemal complex | GO:0000795 | 9.33E-03 |
| Svalbard | synaptonemal structure | GO:0099086 | 9.33E-03 |
| Svalbard | cytoplasmic region | GO:0099568 | 2.10E-02 |
| Svalbard | cell projection | GO:0042995 | 2.78E-02 |
| Svalbard | plasma membrane bounded cell projection | GO:0120025 | 3.55E-02 |
| Svalbard | condensed nuclear chromosome | GO:0000794 | 4.10E-02 |
| France (Pyrenees) | cilium | GO:0005929 | 1.74E-09 |
| France (Pyrenees) | microtubule cytoskeleton | GO:0015630 | 4.08E-06 |
| France (Pyrenees) | cell projection | GO:0042995 | 2.85E-05 |
| France (Pyrenees) | microtubule organizing center | GO:0005815 | 4.46E-05 |
| France (Pyrenees) | plasma membrane bounded cell projection | GO:0120025 | 1.65E-04 |
| France (Pyrenees) | ciliary basal body | GO:0036064 | 6.11E-04 |
| France (Pyrenees) | centrosome | GO:0005813 | 1.33E-03 |
| France (Pyrenees) | ciliary plasm | GO:0097014 | 1.67E-03 |
| France (Pyrenees) | axoneme | GO:0005930 | 3.15E-03 |
| France (Pyrenees) | site of double-strand break | GO:0035861 | 3.82E-03 |
| France (Pyrenees) | ciliary transition fiber | GO:0097539 | 5.28E-03 |

| | | | |
|---|---|---|---|
| France (Pyrenees) | centriole | GO:0005814 | 5.50E-03 |
| France (Pyrenees) | site of DNA damage | GO:0090734 | 8.33E-03 |
| France (Pyrenees) | cytoplasmic region | GO:0099568 | 3.96E-02 |
| Iceland | microtubule organizing center | GO:0005815 | 7.66E-08 |
| Iceland | microtubule cytoskeleton | GO:0015630 | 3.08E-07 |
| Iceland | cilium | GO:0005929 | 4.59E-07 |
| Iceland | centriole | GO:0005814 | 1.40E-05 |
| Iceland | centrosome | GO:0005813 | 5.59E-05 |
| Iceland | non-membrane-bounded organelle | GO:0043228 | 4.72E-04 |
| Iceland | intracellular non-membrane-bounded organelle | GO:0043232 | 4.72E-04 |
| Iceland | ciliary basal body | GO:0036064 | 5.16E-04 |
| Iceland | axoneme | GO:0005930 | 1.50E-03 |
| Iceland | ciliary plasm | GO:0097014 | 2.26E-03 |
| Iceland | site of double-strand break | GO:0035861 | 6.29E-03 |
| Iceland | lateral element | GO:0000800 | 2.12E-02 |
| Iceland | site of DNA damage | GO:0090734 | 3.16E-02 |
| Iceland | plasma membrane bounded cell projection cytoplasm | GO:0032838 | 4.99E-02 |
| E.Greenland | cilium | GO:0005929 | 2.91E-06 |
| E.Greenland | microtubule organizing center | GO:0005815 | 1.67E-05 |
| E.Greenland | microtubule cytoskeleton | GO:0015630 | 3.67E-05 |
| E.Greenland | ciliary basal body | GO:0036064 | 3.10E-04 |
| E.Greenland | ciliary plasm | GO:0097014 | 3.55E-04 |
| E.Greenland | axoneme | GO:0005930 | 6.19E-04 |
| E.Greenland | centrosome | GO:0005813 | 7.05E-04 |
| E.Greenland | site of double-strand break | GO:0035861 | 9.60E-03 |
| E.Greenland | brush border | GO:0005903 | 4.21E-02 |
| W.Greenland | microtubule cytoskeleton | GO:0015630 | 1.89E-09 |
| W.Greenland | microtubule organizing center | GO:0005815 | 3.16E-09 |
| W.Greenland | cilium | GO:0005929 | 1.38E-07 |
| W.Greenland | centrosome | GO:0005813 | 3.46E-06 |
| W.Greenland | ciliary basal body | GO:0036064 | 5.63E-06 |
| W.Greenland | ciliary plasm | GO:0097014 | 3.54E-04 |
| W.Greenland | axoneme | GO:0005930 | 6.96E-04 |
| W.Greenland | centriole | GO:0005814 | 1.30E-03 |
| W.Greenland | secretory granule | GO:0030141 | 5.42E-03 |
| W.Greenland | centriolar satellite | GO:0034451 | 1.21E-02 |

| | | | |
|---|---|---|---|
| W.Greenland | site of double-strand break | GO:0035861 | 1.21E-02 |
| W.Greenland | cell projection | GO:0042995 | 2.35E-02 |
| W.Greenland | cytoskeleton | GO:0005856 | 2.68E-02 |
| W.Greenland | plasma membrane bounded cell projection | GO:0120025 | 3.33E-02 |
| W.Greenland | nuclear chromosome | GO:0000228 | 4.86E-02 |
| France(Alps) | cilium | GO:0005929 | 4.40E-08 |
| France(Alps) | microtubule cytoskeleton | GO:0015630 | 3.98E-06 |
| France(Alps) | microtubule organizing center | GO:0005815 | 1.97E-05 |
| France(Alps) | centriole | GO:0005814 | 6.23E-04 |
| France(Alps) | ciliary basal body | GO:0036064 | 1.17E-03 |
| France(Alps) | ciliary plasm | GO:0097014 | 1.19E-03 |
| France(Alps) | axoneme | GO:0005930 | 2.12E-03 |
| France(Alps) | cell projection | GO:0042995 | 2.67E-03 |
| France(Alps) | centrosome | GO:0005813 | 3.84E-03 |
| France(Alps) | plasma membrane bounded cell projection | GO:0120025 | 7.71E-03 |
| France(Alps) | myosin complex | GO:0016459 | 8.61E-03 |
| France(Alps) | ciliary transition fiber | GO:0097539 | 1.95E-02 |
| France(Alps) | extracellular matrix | GO:0031012 | 2.10E-02 |
| France(Alps) | plasma membrane bounded cell projection cytoplasm | GO:0032838 | 4.43E-02 |
| France(Alps) | cytoplasmic region | GO:0099568 | 4.45E-02 |
| Sweden | cilium | GO:0005929 | 1.11E-07 |
| Sweden | microtubule organizing center | GO:0005815 | 3.07E-06 |
| Sweden | microtubule cytoskeleton | GO:0015630 | 4.31E-06 |
| Sweden | ciliary basal body | GO:0036064 | 1.29E-04 |
| Sweden | secretory granule | GO:0030141 | 1.15E-03 |
| Sweden | centriole | GO:0005814 | 1.54E-03 |
| Sweden | ciliary transition fiber | GO:0097539 | 1.65E-03 |
| Sweden | centrosome | GO:0005813 | 2.65E-03 |
| Sweden | extracellular matrix | GO:0031012 | 2.94E-03 |
| Sweden | axoneme | GO:0005930 | 3.39E-03 |
| Sweden | ciliary plasm | GO:0097014 | 5.14E-03 |
| Sweden | site of double-strand break | GO:0035861 | 6.28E-03 |
| Sweden | collagen-containing extracellular matrix | GO:0062023 | 6.48E-03 |
| Sweden | DNA repair complex | GO:1990391 | 2.38E-02 |
| Sweden | site of DNA damage | GO:0090734 | 4.93E-02 |
| England | microtubule cytoskeleton | GO:0015630 | 8.28E-13 |

| | | | |
|---|---|---|---|
| England | cilium | GO:0005929 | 7.67E-08 |
| England | microtubule organizing center | GO:0005815 | 3.66E-06 |
| England | extracellular matrix | GO:0031012 | 2.10E-04 |
| England | centrosome | GO:0005813 | 2.45E-04 |
| England | collagen-containing extracellular matrix | GO:0062023 | 2.75E-03 |
| England | ciliary plasm | GO:0097014 | 2.97E-03 |
| England | ciliary basal body | GO:0036064 | 5.02E-03 |
| England | axoneme | GO:0005930 | 5.40E-03 |
| England | cytoplasmic region | GO:0099568 | 5.78E-03 |
| England | basement membrane | GO:0005604 | 8.58E-03 |
| England | centriole | GO:0005814 | 1.03E-02 |
| England | cytoskeleton | GO:0005856 | 1.34E-02 |
| England | cell projection | GO:0042995 | 1.42E-02 |
| England | mitotic spindle | GO:0072686 | 1.57E-02 |
| England | microtubule | GO:0005874 | 1.71E-02 |
| England | axonemal dynein complex | GO:0005858 | 2.34E-02 |
| England | spindle | GO:0005819 | 2.36E-02 |
| England | plasma membrane bounded cell projection | GO:0120025 | 2.86E-02 |
| England | intracellular non-membrane-bounded organelle | GO:0043232 | 3.31E-02 |
| England | non-membrane-bounded organelle | GO:0043228 | 3.31E-02 |
| Canada (Newfoundland) | cilium | GO:0005929 | 1.45E-09 |
| Canada (Newfoundland) | microtubule cytoskeleton | GO:0015630 | 1.50E-09 |
| Canada (Newfoundland) | microtubule organizing center | GO:0005815 | 8.52E-08 |
| Canada (Newfoundland) | centrosome | GO:0005813 | 2.30E-05 |
| Canada (Newfoundland) | ciliary basal body | GO:0036064 | 2.68E-04 |
| Canada (Newfoundland) | cell projection | GO:0042995 | 6.76E-04 |
| Canada (Newfoundland) | plasma membrane bounded cell projection | GO:0120025 | 1.58E-03 |
| Canada (Newfoundland) | spindle | GO:0005819 | 3.40E-03 |
| Canada (Newfoundland) | ciliary plasm | GO:0097014 | 1.36E-02 |
| Canada (Newfoundland) | centriole | GO:0005814 | 2.20E-02 |
| Canada (Newfoundland) | axoneme | GO:0005930 | 2.24E-02 |
| Canada (Newfoundland) | site of double-strand break | GO:0035861 | 2.28E-02 |
| Canada (Newfoundland) | motile cilium | GO:0031514 | 2.85E-02 |
| Canada (Newfoundland) | extracellular matrix | GO:0031012 | 3.51E-02 |
| Canada (Newfoundland) | cytoskeleton | GO:0005856 | 4.74E-02 |
| Norway | microtubule cytoskeleton | GO:0015630 | 9.71E-08 |

| | | | |
|---|---|---|---|
| Norway | cilium | GO:0005929 | 3.65E-06 |
| Norway | microtubule organizing center | GO:0005815 | 5.03E-05 |
| Norway | centriole | GO:0005814 | 3.85E-04 |
| Norway | ciliary basal body | GO:0036064 | 1.47E-03 |
| Norway | centrosome | GO:0005813 | 1.77E-03 |
| Norway | site of double-strand break | GO:0035861 | 1.27E-02 |
| Norway | ciliary plasm | GO:0097014 | 1.68E-02 |
| Norway | axoneme | GO:0005930 | 2.79E-02 |
| Norway | spindle | GO:0005819 | 3.18E-02 |
| Norway | ciliary transition fiber | GO:0097539 | 3.46E-02 |
| Norway | cell projection | GO:0042995 | 3.71E-02 |

**b)**
**GO:Biological process**

| Pop | term_name | term_id | adjusted_p_value |
|---|---|---|---|
| Svalbard | cilium organization | GO:0044782 | 4.20E-06 |
| Svalbard | cilium assembly | GO:0060271 | 8.88E-05 |
| Svalbard | microtubule-based process | GO:0007017 | 1.97E-04 |
| Svalbard | microtubule-based movement | GO:0007018 | 3.01E-03 |
| Svalbard | plasma membrane bounded cell projection assembly | GO:0120031 | 5.06E-03 |
| Svalbard | cell projection assembly | GO:0030031 | 8.85E-03 |
| Svalbard | organelle assembly | GO:0070925 | 1.57E-02 |
| France (Pyrenees) | cilium organization | GO:0044782 | 3.72E-11 |
| France (Pyrenees) | cilium assembly | GO:0060271 | 2.98E-09 |
| France (Pyrenees) | organelle assembly | GO:0070925 | 1.37E-07 |
| France (Pyrenees) | microtubule-based process | GO:0007017 | 2.06E-07 |
| France (Pyrenees) | plasma membrane bounded cell projection organization | GO:0120036 | 1.41E-06 |
| France (Pyrenees) | cell projection organization | GO:0030030 | 1.48E-06 |
| France (Pyrenees) | microtubule cytoskeleton organization | GO:0000226 | 1.10E-05 |
| France (Pyrenees) | cell projection assembly | GO:0030031 | 1.49E-05 |
| France (Pyrenees) | plasma membrane bounded cell projection assembly | GO:0120031 | 1.83E-05 |
| France (Pyrenees) | organelle organization | GO:0006996 | 1.65E-03 |
| France (Pyrenees) | microtubule-based movement | GO:0007018 | 5.48E-03 |
| France (Pyrenees) | cytoskeleton organization | GO:0007010 | 7.61E-03 |

| | | | |
|---|---|---|---|
| France (Pyrenees) | double-strand break repair | GO:0006302 | 1.46E-02 |
| France (Pyrenees) | cell cycle process | GO:0022402 | 1.99E-02 |
| Iceland | cilium assembly | GO:0060271 | 1.68E-07 |
| Iceland | cilium organization | GO:0044782 | 3.13E-07 |
| Iceland | organelle assembly | GO:0070925 | 1.17E-05 |
| Iceland | plasma membrane bounded cell projection assembly | GO:0120031 | 4.09E-05 |
| Iceland | cell projection assembly | GO:0030031 | 8.10E-05 |
| Iceland | microtubule-based process | GO:0007017 | 3.26E-03 |
| Iceland | microtubule organizing center organization | GO:0031023 | 3.29E-03 |
| Iceland | centrosome cycle | GO:0007098 | 1.38E-02 |
| Iceland | microtubule-based movement | GO:0007018 | 4.90E-02 |
| E.Greenland | cilium organization | GO:0044782 | 9.67E-15 |
| E.Greenland | cilium assembly | GO:0060271 | 1.97E-14 |
| E.Greenland | plasma membrane bounded cell projection assembly | GO:0120031 | 7.01E-10 |
| E.Greenland | microtubule-based process | GO:0007017 | 7.21E-10 |
| E.Greenland | cell projection assembly | GO:0030031 | 8.13E-10 |
| E.Greenland | microtubule-based movement | GO:0007018 | 7.19E-09 |
| E.Greenland | organelle assembly | GO:0070925 | 1.62E-08 |
| E.Greenland | DNA repair | GO:0006281 | 1.78E-04 |
| E.Greenland | microtubule cytoskeleton organization | GO:0000226 | 5.19E-04 |
| E.Greenland | cilium movement | GO:0003341 | 3.81E-03 |
| E.Greenland | nuclear chromosome segregation | GO:0098813 | 4.81E-03 |
| E.Greenland | double-strand break repair | GO:0006302 | 5.28E-03 |
| E.Greenland | meiotic chromosome segregation | GO:0045132 | 6.91E-03 |
| E.Greenland | intraciliary retrograde transport | GO:0035721 | 8.61E-03 |
| E.Greenland | axoneme assembly | GO:0035082 | 1.20E-02 |
| E.Greenland | chromosome segregation | GO:0007059 | 1.36E-02 |
| E.Greenland | DNA recombination | GO:0006310 | 1.37E-02 |
| E.Greenland | microtubule-based transport | GO:0099111 | 1.42E-02 |
| E.Greenland | plasma membrane bounded cell projection organization | GO:0120036 | 3.15E-02 |
| E.Greenland | homophilic cell adhesion via plasma membrane adhesion molecules | GO:0007156 | 3.43E-02 |
| E.Greenland | cell projection organization | GO:0030030 | 3.71E-02 |
| W.Greenland | cilium organization | GO:0044782 | 2.52E-11 |
| W.Greenland | cilium assembly | GO:0060271 | 1.16E-10 |
| W.Greenland | microtubule-based process | GO:0007017 | 1.08E-08 |
| W.Greenland | organelle assembly | GO:0070925 | 1.65E-07 |

| | | | |
|---|---|---|---|
| W.Greenland | cell projection assembly | GO:0030031 | 8.79E-07 |
| W.Greenland | plasma membrane bounded cell projection assembly | GO:0120031 | 1.02E-06 |
| W.Greenland | double-strand break repair | GO:0006302 | 4.55E-05 |
| W.Greenland | microtubule-based movement | GO:0007018 | 6.55E-05 |
| W.Greenland | microtubule cytoskeleton organization | GO:0000226 | 9.18E-05 |
| W.Greenland | organelle organization | GO:0006996 | 3.36E-03 |
| W.Greenland | plasma membrane bounded cell projection organization | GO:0120036 | 4.36E-03 |
| W.Greenland | cell projection organization | GO:0030030 | 4.51E-03 |
| W.Greenland | cell cycle process | GO:0022402 | 5.91E-03 |
| W.Greenland | double-strand break repair via homologous recombination | GO:0000724 | 8.23E-03 |
| W.Greenland | recombinational repair | GO:0000725 | 9.92E-03 |
| W.Greenland | DNA repair | GO:0006281 | 1.05E-02 |
| W.Greenland | DNA recombination | GO:0006310 | 2.04E-02 |
| W.Greenland | DNA replication | GO:0006260 | 3.15E-02 |
| W.Greenland | cytoskeleton organization | GO:0007010 | 3.23E-02 |
| France(Alps) | cilium organization | GO:0044782 | 2.23E-11 |
| France(Alps) | cilium assembly | GO:0060271 | 8.91E-10 |
| France(Alps) | microtubule-based process | GO:0007017 | 1.46E-09 |
| France(Alps) | organelle assembly | GO:0070925 | 1.85E-07 |
| France(Alps) | cell projection assembly | GO:0030031 | 1.41E-06 |
| France(Alps) | plasma membrane bounded cell projection assembly | GO:0120031 | 1.44E-06 |
| France(Alps) | microtubule-based movement | GO:0007018 | 1.66E-06 |
| France(Alps) | microtubule cytoskeleton organization | GO:0000226 | 4.28E-06 |
| France(Alps) | axoneme assembly | GO:0035082 | 7.13E-05 |
| France(Alps) | double-strand break repair | GO:0006302 | 3.40E-04 |
| France(Alps) | organelle organization | GO:0006996 | 6.17E-04 |
| France(Alps) | meiotic chromosome segregation | GO:0045132 | 1.21E-03 |
| France(Alps) | cell cycle process | GO:0022402 | 1.70E-03 |
| France(Alps) | DNA repair | GO:0006281 | 2.73E-03 |
| France(Alps) | microtubule bundle formation | GO:0001578 | 2.77E-03 |
| France(Alps) | cilium movement | GO:0003341 | 5.76E-03 |
| France(Alps) | cytoskeleton organization | GO:0007010 | 6.68E-03 |
| France(Alps) | meiotic nuclear division | GO:0140013 | 8.52E-03 |
| France(Alps) | male meiotic nuclear division | GO:0007140 | 8.64E-03 |
| France(Alps) | organelle fission | GO:0048285 | 9.17E-03 |
| France(Alps) | meiotic cell cycle process | GO:1903046 | 9.27E-03 |

| | | | |
|---|---|---|---|
| France(Alps) | chromosome segregation | GO:0007059 | 9.55E-03 |
| France(Alps) | DNA double-strand break processing | GO:0000729 | 2.51E-02 |
| France(Alps) | meiotic cell cycle | GO:0051321 | 3.41E-02 |
| France(Alps) | microtubule organizing center organization | GO:0031023 | 3.76E-02 |
| France(Alps) | DNA recombination | GO:0006310 | 4.28E-02 |
| Sweden | cilium assembly | GO:0060271 | 4.99E-05 |
| Sweden | cilium organization | GO:0044782 | 8.00E-05 |
| Sweden | microtubule-based process | GO:0007017 | 9.62E-04 |
| Sweden | organelle assembly | GO:0070925 | 3.71E-03 |
| Sweden | DNA replication | GO:0006260 | 1.08E-02 |
| Sweden | microtubule-based movement | GO:0007018 | 1.33E-02 |
| Sweden | double-strand break repair | GO:0006302 | 1.39E-02 |
| Sweden | cilium movement | GO:0003341 | 1.48E-02 |
| Sweden | plasma membrane bounded cell projection assembly | GO:0120031 | 2.08E-02 |
| Sweden | microtubule cytoskeleton organization | GO:0000226 | 2.29E-02 |
| Sweden | cell cycle process | GO:0022402 | 3.26E-02 |
| Sweden | cell projection assembly | GO:0030031 | 3.52E-02 |
| Sweden | DNA-dependent DNA replication | GO:0006261 | 3.80E-02 |
| England | microtubule-based process | GO:0007017 | 2.99E-11 |
| England | cilium organization | GO:0044782 | 1.52E-10 |
| England | cilium assembly | GO:0060271 | 1.86E-09 |
| England | cellular response to DNA damage stimulus | GO:0006974 | 2.62E-07 |
| England | double-strand break repair | GO:0006302 | 7.92E-07 |
| England | microtubule-based movement | GO:0007018 | 1.01E-06 |
| England | organelle assembly | GO:0070925 | 2.31E-06 |
| England | DNA repair | GO:0006281 | 2.82E-06 |
| England | microtubule cytoskeleton organization | GO:0000226 | 3.52E-06 |
| England | cell projection assembly | GO:0030031 | 2.40E-05 |
| England | plasma membrane bounded cell projection assembly | GO:0120031 | 2.58E-05 |
| England | DNA recombination | GO:0006310 | 9.41E-05 |
| England | regulation of DNA replication | GO:0006275 | 1.28E-03 |
| England | double-strand break repair via homologous recombination | GO:0000724 | 1.42E-03 |
| England | recombinational repair | GO:0000725 | 1.77E-03 |
| England | microtubule-based transport | GO:0099111 | 6.44E-03 |
| England | DNA replication | GO:0006260 | 7.00E-03 |
| England | organelle organization | GO:0006996 | 1.14E-02 |

| | | | |
|---|---|---|---|
| England | mitotic cell cycle checkpoint | GO:0007093 | 1.48E-02 |
| England | DNA-dependent DNA replication | GO:0006261 | 2.95E-02 |
| England | negative regulation of DNA metabolic process | GO:0051053 | 2.97E-02 |
| England | intraciliary retrograde transport | GO:0035721 | 4.27E-02 |
| Canada (Newfoundland) | microtubule cytoskeleton organization | GO:0000226 | 3.08E-11 |
| Canada (Newfoundland) | microtubule-based process | GO:0007017 | 1.09E-10 |
| Canada (Newfoundland) | cilium assembly | GO:0060271 | 4.97E-09 |
| Canada (Newfoundland) | cilium organization | GO:0044782 | 2.12E-08 |
| Canada (Newfoundland) | organelle assembly | GO:0070925 | 3.95E-06 |
| Canada (Newfoundland) | cell cycle process | GO:0022402 | 6.71E-06 |
| Canada (Newfoundland) | microtubule bundle formation | GO:0001578 | 4.44E-05 |
| Canada (Newfoundland) | axoneme assembly | GO:0035082 | 5.52E-05 |
| Canada (Newfoundland) | plasma membrane bounded cell projection assembly | GO:0120031 | 8.17E-05 |
| Canada (Newfoundland) | cell projection assembly | GO:0030031 | 8.35E-05 |
| Canada (Newfoundland) | double-strand break repair | GO:0006302 | 4.73E-04 |
| Canada (Newfoundland) | microtubule organizing center organization | GO:0031023 | 6.45E-04 |
| Canada (Newfoundland) | cell cycle | GO:0007049 | 8.98E-04 |
| Canada (Newfoundland) | cytoskeleton organization | GO:0007010 | 1.32E-03 |
| Canada (Newfoundland) | centrosome duplication | GO:0051298 | 4.26E-03 |
| Canada (Newfoundland) | centrosome cycle | GO:0007098 | 4.62E-03 |
| Canada (Newfoundland) | microtubule-based movement | GO:0007018 | 5.03E-03 |
| Canada (Newfoundland) | DNA repair | GO:0006281 | 6.03E-03 |
| Canada (Newfoundland) | nuclear division | GO:0000280 | 8.49E-03 |
| Canada (Newfoundland) | double-strand break repair via homologous recombination | GO:0000724 | 1.81E-02 |
| Canada (Newfoundland) | mitotic cell cycle | GO:0000278 | 1.95E-02 |
| Canada (Newfoundland) | recombinational repair | GO:0000725 | 2.22E-02 |
| Canada (Newfoundland) | DNA recombination | GO:0006310 | 2.51E-02 |
| Canada (Newfoundland) | sperm axoneme assembly | GO:0007288 | 3.99E-02 |
| Canada (Newfoundland) | chromosome segregation | GO:0007059 | 4.93E-02 |
| Norway | cilium organization | GO:0044782 | 3.29E-11 |
| Norway | cilium assembly | GO:0060271 | 2.64E-10 |
| Norway | microtubule-based process | GO:0007017 | 2.26E-07 |
| Norway | plasma membrane bounded cell projection assembly | GO:0120031 | 3.30E-07 |
| Norway | cell projection assembly | GO:0030031 | 3.60E-07 |
| Norway | organelle assembly | GO:0070925 | 4.85E-07 |
| Norway | double-strand break repair | GO:0006302 | 2.58E-05 |

| | | | |
|---|---|---|---|
| Norway | microtubule cytoskeleton organization | GO:0000226 | 2.79E-05 |
| Norway | microtubule-based movement | GO:0007018 | 2.54E-04 |
| Norway | DNA recombination | GO:0006310 | 2.97E-04 |
| Norway | regulation of DNA replication | GO:0006275 | 9.31E-04 |
| Norway | DNA repair | GO:0006281 | 1.03E-03 |
| Norway | microtubule bundle formation | GO:0001578 | 1.22E-03 |
| Norway | double-strand break repair via homologous recombination | GO:0000724 | 7.29E-03 |
| Norway | recombinational repair | GO:0000725 | 9.06E-03 |
| Norway | axoneme assembly | GO:0035082 | 1.29E-02 |
| Norway | cilium movement | GO:0003341 | 1.38E-02 |
| Norway | cellular response to DNA damage stimulus | GO:0006974 | 2.60E-02 |
| Norway | DNA replication | GO:0006260 | 2.94E-02 |

**GO:Molecular function**

| Pop | term_name | term_id | adjusted_p_value |
|---|---|---|---|
| France (Pyrenees) | motor activity | GO:0003774 | 1.67E-05 |
| France (Pyrenees) | cytoskeletal protein binding | GO:0008092 | 1.78E-04 |
| France (Pyrenees) | microtubule motor activity | GO:0003777 | 2.31E-03 |
| France (Pyrenees) | tubulin binding | GO:0015631 | 2.82E-03 |
| France (Pyrenees) | ATPase activity | GO:0016887 | 4.84E-03 |
| France (Pyrenees) | nucleoside-triphosphatase activity | GO:0017111 | 1.18E-02 |
| France (Pyrenees) | pyrophosphatase activity | GO:0016462 | 1.28E-02 |
| France (Pyrenees) | microtubule binding | GO:0008017 | 1.48E-02 |
| France (Pyrenees) | hydrolase activity, acting on acid anhydrides, in phosphorus-containing anhydrides | GO:0016818 | 1.51E-02 |
| France (Pyrenees) | hydrolase activity, acting on acid anhydrides | GO:0016817 | 1.67E-02 |
| Iceland | ATPase activity | GO:0016887 | 3.87E-02 |
| E.Greenland | motor activity | GO:0003774 | 7.51E-03 |
| E.Greenland | microtubule motor activity | GO:0003777 | 2.44E-02 |
| E.Greenland | calcium ion binding | GO:0005509 | 3.49E-02 |
| W.Greenland | motor activity | GO:0003774 | 2.99E-03 |
| W.Greenland | microtubule motor activity | GO:0003777 | 5.72E-03 |
| W.Greenland | oxidoreductase activity | GO:0016491 | 2.35E-02 |
| W.Greenland | catalytic activity, acting on DNA | GO:0140097 | 3.10E-02 |
| W.Greenland | ATPase activity | GO:0016887 | 4.91E-02 |
| France(Alps) | ATPase activity | GO:0016887 | 1.45E-04 |
| France(Alps) | motor activity | GO:0003774 | 2.07E-03 |

| | | | |
|---|---|---|---|
| France(Alps) | ATP binding | GO:0005524 | 2.07E-03 |
| France(Alps) | adenyl nucleotide binding | GO:0030554 | 2.32E-03 |
| France(Alps) | adenyl ribonucleotide binding | GO:0032559 | 4.09E-03 |
| France(Alps) | nucleoside-triphosphatase activity | GO:0017111 | 1.22E-02 |
| France(Alps) | ion binding | GO:0043167 | 1.90E-02 |
| France(Alps) | pyrophosphatase activity | GO:0016462 | 1.96E-02 |
| France(Alps) | hydrolase activity, acting on acid anhydrides, in phosphorus-containing anhydrides | GO:0016818 | 2.34E-02 |
| France(Alps) | hydrolase activity, acting on acid anhydrides | GO:0016817 | 2.64E-02 |
| Sweden | ATPase activity | GO:0016887 | 3.76E-04 |
| England | ATPase activity | GO:0016887 | 1.08E-07 |
| England | catalytic activity, acting on DNA | GO:0140097 | 1.16E-03 |
| England | microtubule motor activity | GO:0003777 | 2.45E-03 |
| England | helicase activity | GO:0004386 | 3.70E-03 |
| England | DNA helicase activity | GO:0003678 | 3.70E-03 |
| England | tubulin binding | GO:0015631 | 4.73E-03 |
| England | motor activity | GO:0003774 | 1.04E-02 |
| England | microtubule binding | GO:0008017 | 2.01E-02 |
| Canada (Newfoundland) | tubulin binding | GO:0015631 | 5.51E-04 |
| Canada (Newfoundland) | microtubule binding | GO:0008017 | 7.96E-04 |
| Canada (Newfoundland) | ATPase activity | GO:0016887 | 1.51E-03 |
| Canada (Newfoundland) | iron ion binding | GO:0005506 | 4.26E-02 |
| Norway | ATPase activity | GO:0016887 | 1.68E-05 |
| Norway | motor activity | GO:0003774 | 8.94E-05 |
| Norway | ATP binding | GO:0005524 | 5.34E-03 |
| Norway | adenyl nucleotide binding | GO:0030554 | 7.37E-03 |
| Norway | nucleoside-triphosphatase activity | GO:0017111 | 7.77E-03 |
| Norway | pyrophosphatase activity | GO:0016462 | 9.58E-03 |
| Norway | hydrolase activity, acting on acid anhydrides, in phosphorus-containing anhydrides | GO:0016818 | 1.17E-02 |
| Norway | adenyl ribonucleotide binding | GO:0032559 | 1.18E-02 |
| Norway | tubulin binding | GO:0015631 | 1.21E-02 |
| Norway | hydrolase activity, acting on acid anhydrides | GO:0016817 | 1.33E-02 |
| Norway | cytoskeletal protein binding | GO:0008092 | 2.38E-02 |

**GO:Cellular component**

| Pop | term_name | term_id | adjusted_p_value |
|---|---|---|---|
| Svalbard | cilium | GO:0005929 | 2.42E-06 |

| | | | |
|---|---|---|---|
| Svalbard | microtubule cytoskeleton | GO:0015630 | 3.96E-04 |
| Svalbard | ciliary transition fiber | GO:0097539 | 1.00E-03 |
| Svalbard | axoneme | GO:0005930 | 2.33E-03 |
| Svalbard | ciliary plasm | GO:0097014 | 2.83E-03 |
| Svalbard | cell projection | GO:0042995 | 3.02E-03 |
| Svalbard | ciliary base | GO:0097546 | 3.42E-03 |
| Svalbard | ciliary basal body | GO:0036064 | 4.04E-03 |
| Svalbard | plasma membrane bounded cell projection | GO:0120025 | 8.45E-03 |
| Svalbard | lateral element | GO:0000800 | 2.32E-02 |
| Svalbard | microtubule organizing center | GO:0005815 | 2.78E-02 |
| Svalbard | centriole | GO:0005814 | 2.87E-02 |
| Svalbard | cytoskeleton | GO:0005856 | 4.70E-02 |
| Svalbard | cytoplasmic region | GO:0099568 | 4.87E-02 |
| France (Pyrenees) | microtubule cytoskeleton | GO:0015630 | 1.56E-12 |
| France (Pyrenees) | cilium | GO:0005929 | 7.23E-09 |
| France (Pyrenees) | intracellular non-membrane-bounded organelle | GO:0043232 | 1.45E-08 |
| France (Pyrenees) | non-membrane-bounded organelle | GO:0043228 | 1.45E-08 |
| France (Pyrenees) | cytoskeleton | GO:0005856 | 1.72E-08 |
| France (Pyrenees) | microtubule organizing center | GO:0005815 | 2.47E-07 |
| France (Pyrenees) | centrosome | GO:0005813 | 3.93E-06 |
| France (Pyrenees) | plasma membrane bounded cell projection | GO:0120025 | 1.12E-04 |
| France (Pyrenees) | cell projection | GO:0042995 | 1.43E-04 |
| France (Pyrenees) | centriole | GO:0005814 | 1.62E-04 |
| France (Pyrenees) | ciliary basal body | GO:0036064 | 1.05E-03 |
| France (Pyrenees) | spindle | GO:0005819 | 3.62E-03 |
| France (Pyrenees) | ciliary plasm | GO:0097014 | 4.93E-03 |
| France (Pyrenees) | axonemal dynein complex | GO:0005858 | 1.30E-02 |
| France (Pyrenees) | plasma membrane bounded cell projection cytoplasm | GO:0032838 | 1.43E-02 |
| France (Pyrenees) | cytoplasmic region | GO:0099568 | 1.59E-02 |
| France (Pyrenees) | axoneme | GO:0005930 | 1.66E-02 |
| France (Pyrenees) | dynein complex | GO:0030286 | 1.70E-02 |
| Iceland | microtubule cytoskeleton | GO:0015630 | 5.23E-05 |
| Iceland | laminin complex | GO:0043256 | 2.23E-04 |
| Iceland | cilium | GO:0005929 | 3.85E-04 |
| Iceland | microtubule organizing center | GO:0005815 | 1.07E-03 |
| Iceland | centrosome | GO:0005813 | 4.15E-03 |

| | | | |
|---|---|---|---|
| Iceland | basement membrane | GO:0005604 | 1.11E-02 |
| Iceland | centriole | GO:0005814 | 2.54E-02 |
| Iceland | intracellular non-membrane-bounded organelle | GO:0043232 | 4.02E-02 |
| Iceland | non-membrane-bounded organelle | GO:0043228 | 4.02E-02 |
| E.Greenland | cilium | GO:0005929 | 2.29E-10 |
| E.Greenland | microtubule cytoskeleton | GO:0015630 | 1.46E-09 |
| E.Greenland | microtubule organizing center | GO:0005815 | 1.20E-08 |
| E.Greenland | ciliary plasm | GO:0097014 | 7.73E-07 |
| E.Greenland | axoneme | GO:0005930 | 2.91E-06 |
| E.Greenland | ciliary basal body | GO:0036064 | 4.21E-06 |
| E.Greenland | centrosome | GO:0005813 | 5.90E-06 |
| E.Greenland | cytoskeleton | GO:0005856 | 9.14E-06 |
| E.Greenland | cytoplasmic region | GO:0099568 | 4.47E-05 |
| E.Greenland | plasma membrane bounded cell projection cytoplasm | GO:0032838 | 3.40E-04 |
| E.Greenland | dynein complex | GO:0030286 | 1.15E-03 |
| E.Greenland | centriole | GO:0005814 | 1.50E-03 |
| E.Greenland | non-membrane-bounded organelle | GO:0043228 | 3.01E-03 |
| E.Greenland | intracellular non-membrane-bounded organelle | GO:0043232 | 3.01E-03 |
| E.Greenland | cell projection | GO:0042995 | 3.60E-03 |
| E.Greenland | Fanconi anaemia nuclear complex | GO:0043240 | 6.63E-03 |
| E.Greenland | plasma membrane bounded cell projection | GO:0120025 | 1.19E-02 |
| E.Greenland | ciliary transition fiber | GO:0097539 | 1.84E-02 |
| W.Greenland | microtubule organizing center | GO:0005815 | 2.05E-10 |
| W.Greenland | microtubule cytoskeleton | GO:0015630 | 2.31E-09 |
| W.Greenland | centrosome | GO:0005813 | 1.32E-08 |
| W.Greenland | cilium | GO:0005929 | 9.35E-07 |
| W.Greenland | ciliary basal body | GO:0036064 | 1.71E-05 |
| W.Greenland | centriole | GO:0005814 | 1.08E-04 |
| W.Greenland | axoneme | GO:0005930 | 5.51E-04 |
| W.Greenland | ciliary plasm | GO:0097014 | 7.00E-04 |
| W.Greenland | cytoskeleton | GO:0005856 | 8.74E-04 |
| W.Greenland | cell projection | GO:0042995 | 1.01E-03 |
| W.Greenland | plasma membrane bounded cell projection | GO:0120025 | 3.45E-03 |
| W.Greenland | ciliary transition fiber | GO:0097539 | 4.41E-03 |
| W.Greenland | intracellular non-membrane-bounded organelle | GO:0043232 | 1.36E-02 |
| W.Greenland | non-membrane-bounded organelle | GO:0043228 | 1.36E-02 |

| | | | |
|---|---|---|---|
| W.Greenland | cytoplasmic region | GO:0099568 | 1.56E-02 |
| W.Greenland | ciliary base | GO:0097546 | 2.10E-02 |
| W.Greenland | axonemal dynein complex | GO:0005858 | 2.12E-02 |
| W.Greenland | dynein complex | GO:0030286 | 3.60E-02 |
| W.Greenland | plasma membrane bounded cell projection cytoplasm | GO:0032838 | 3.76E-02 |
| France(Alps) | microtubule cytoskeleton | GO:0015630 | 2.19E-13 |
| France(Alps) | cilium | GO:0005929 | 1.78E-10 |
| France(Alps) | microtubule organizing center | GO:0005815 | 1.91E-09 |
| France(Alps) | cytoskeleton | GO:0005856 | 7.88E-08 |
| France(Alps) | centrosome | GO:0005813 | 9.54E-07 |
| France(Alps) | ciliary plasm | GO:0097014 | 4.58E-06 |
| France(Alps) | dynein complex | GO:0030286 | 1.08E-05 |
| France(Alps) | non-membrane-bounded organelle | GO:0043228 | 1.49E-05 |
| France(Alps) | intracellular non-membrane-bounded organelle | GO:0043232 | 1.49E-05 |
| France(Alps) | axoneme | GO:0005930 | 1.72E-05 |
| France(Alps) | basement membrane | GO:0005604 | 2.34E-05 |
| France(Alps) | collagen-containing extracellular matrix | GO:0062023 | 5.28E-05 |
| France(Alps) | plasma membrane bounded cell projection cytoplasm | GO:0032838 | 7.60E-05 |
| France(Alps) | ciliary basal body | GO:0036064 | 1.53E-04 |
| France(Alps) | cytoplasmic region | GO:0099568 | 3.28E-04 |
| France(Alps) | ciliary transition fiber | GO:0097539 | 5.11E-04 |
| France(Alps) | microtubule associated complex | GO:0005875 | 1.35E-03 |
| France(Alps) | centriole | GO:0005814 | 1.70E-03 |
| France(Alps) | cell projection | GO:0042995 | 6.72E-03 |
| France(Alps) | plasma membrane bounded cell projection | GO:0120025 | 1.62E-02 |
| France(Alps) | extracellular matrix | GO:0031012 | 3.90E-02 |
| Sweden | cilium | GO:0005929 | 8.08E-13 |
| Sweden | microtubule cytoskeleton | GO:0015630 | 4.92E-09 |
| Sweden | microtubule organizing center | GO:0005815 | 7.25E-09 |
| Sweden | centrosome | GO:0005813 | 7.16E-07 |
| Sweden | ciliary transition fiber | GO:0097539 | 3.49E-05 |
| Sweden | centriole | GO:0005814 | 1.20E-04 |
| Sweden | ciliary basal body | GO:0036064 | 1.31E-04 |
| Sweden | cell projection | GO:0042995 | 2.84E-04 |
| Sweden | plasma membrane bounded cell projection | GO:0120025 | 4.56E-04 |
| Sweden | cytoskeleton | GO:0005856 | 1.15E-03 |

| | | | |
|---|---|---|---|
| Sweden | DNA repair complex | GO:1990391 | 3.58E-03 |
| Sweden | axoneme | GO:0005930 | 3.70E-03 |
| Sweden | intracellular non-membrane-bounded organelle | GO:0043232 | 4.18E-03 |
| Sweden | non-membrane-bounded organelle | GO:0043228 | 4.18E-03 |
| Sweden | secretory granule | GO:0030141 | 4.24E-03 |
| Sweden | ciliary plasm | GO:0097014 | 4.49E-03 |
| Sweden | secretory vesicle | GO:0099503 | 2.27E-02 |
| Sweden | plasma membrane bounded cell projection cytoplasm | GO:0032838 | 3.49E-02 |
| England | microtubule cytoskeleton | GO:0015630 | 1.40E-13 |
| England | cilium | GO:0005929 | 4.26E-09 |
| England | microtubule organizing center | GO:0005815 | 1.88E-08 |
| England | cytoplasmic region | GO:0099568 | 2.19E-08 |
| England | centrosome | GO:0005813 | 1.05E-06 |
| England | plasma membrane bounded cell projection cytoplasm | GO:0032838 | 1.20E-06 |
| England | axoneme | GO:0005930 | 5.94E-06 |
| England | ciliary plasm | GO:0097014 | 8.11E-06 |
| England | non-membrane-bounded organelle | GO:0043228 | 2.61E-04 |
| England | intracellular non-membrane-bounded organelle | GO:0043232 | 2.61E-04 |
| England | ciliary basal body | GO:0036064 | 6.25E-04 |
| England | dynein complex | GO:0030286 | 1.03E-03 |
| England | cytoskeleton | GO:0005856 | 1.10E-03 |
| England | axonemal dynein complex | GO:0005858 | 2.63E-03 |
| England | microtubule | GO:0005874 | 3.01E-03 |
| England | non-motile cilium | GO:0097730 | 5.22E-03 |
| England | microtubule associated complex | GO:0005875 | 6.27E-03 |
| England | centriole | GO:0005814 | 1.04E-02 |
| England | spindle | GO:0005819 | 1.16E-02 |
| England | cell projection | GO:0042995 | 2.01E-02 |
| England | RNA polymerase III transcription regulator complex | GO:0090576 | 2.55E-02 |
| England | plasma membrane bounded cell projection | GO:0120025 | 3.37E-02 |
| Canada (Newfoundland) | microtubule cytoskeleton | GO:0015630 | 5.70E-11 |
| Canada (Newfoundland) | cilium | GO:0005929 | 6.36E-08 |
| Canada (Newfoundland) | ciliary basal body | GO:0036064 | 1.47E-07 |
| Canada (Newfoundland) | microtubule organizing center | GO:0005815 | 2.66E-06 |
| Canada (Newfoundland) | cytoskeleton | GO:0005856 | 1.04E-05 |
| Canada (Newfoundland) | cytoplasmic region | GO:0099568 | 2.91E-05 |

| | | | |
|---|---|---|---|
| Canada (Newfoundland) | axoneme | GO:0005930 | 2.37E-04 |
| Canada (Newfoundland) | ciliary plasm | GO:0097014 | 3.13E-04 |
| Canada (Newfoundland) | basement membrane | GO:0005604 | 3.13E-04 |
| Canada (Newfoundland) | cell projection | GO:0042995 | 4.19E-04 |
| Canada (Newfoundland) | plasma membrane bounded cell projection | GO:0120025 | 6.24E-04 |
| Canada (Newfoundland) | spindle | GO:0005819 | 8.40E-04 |
| Canada (Newfoundland) | intracellular non-membrane-bounded organelle | GO:0043232 | 9.38E-04 |
| Canada (Newfoundland) | non-membrane-bounded organelle | GO:0043228 | 9.38E-04 |
| Canada (Newfoundland) | collagen-containing extracellular matrix | GO:0062023 | 1.87E-03 |
| Canada (Newfoundland) | centrosome | GO:0005813 | 2.02E-03 |
| Canada (Newfoundland) | microtubule | GO:0005874 | 3.04E-03 |
| Canada (Newfoundland) | plasma membrane bounded cell projection cytoplasm | GO:0032838 | 1.02E-02 |
| Canada (Newfoundland) | extracellular matrix | GO:0031012 | 3.53E-02 |
| Norway | microtubule cytoskeleton | GO:0015630 | 6.19E-10 |
| Norway | cilium | GO:0005929 | 1.05E-08 |
| Norway | ciliary basal body | GO:0036064 | 1.15E-08 |
| Norway | microtubule organizing center | GO:0005815 | 3.10E-07 |
| Norway | centrosome | GO:0005813 | 5.75E-05 |
| Norway | cytoplasmic region | GO:0099568 | 1.50E-04 |
| Norway | cytoskeleton | GO:0005856 | 2.02E-04 |
| Norway | cell projection | GO:0042995 | 2.10E-04 |
| Norway | centriole | GO:0005814 | 3.76E-04 |
| Norway | dynein complex | GO:0030286 | 1.14E-03 |
| Norway | plasma membrane bounded cell projection | GO:0120025 | 1.25E-03 |
| Norway | axoneme | GO:0005930 | 1.30E-03 |
| Norway | ciliary plasm | GO:0097014 | 1.67E-03 |
| Norway | myosin complex | GO:0016459 | 2.97E-03 |
| Norway | plasma membrane bounded cell projection cytoplasm | GO:0032838 | 4.06E-03 |
| Norway | axonemal dynein complex | GO:0005858 | 6.66E-03 |
| Norway | intracellular non-membrane-bounded organelle | GO:0043232 | 8.94E-03 |
| Norway | non-membrane-bounded organelle | GO:0043228 | 8.94E-03 |
| Norway | microtubule associated complex | GO:0005875 | 1.56E-02 |
| Norway | pericentriolar material | GO:0000242 | 1.79E-02 |
| Norway | basement membrane | GO:0005604 | 2.28E-02 |

**c)**

**GO:Molecular function**

| Pop | term_name | term_id | adjusted_p_value |
|---|---|---|---|
| E.Greenland | guanyl-nucleotide exchange factor activity | GO:0005085 | 3.08E-02 |

**GO:Cellular component**

| Pop | term_name | term_id | adjusted_p_value |
|---|---|---|---|
| France (Pyrenees) | TORC1 complex | GO:0031931 | 7.23E-03 |
| Sweden | nuclear pore | GO:0005643 | 4.76E-02 |
| Canada (Newfoundland) | stereocilia ankle link complex | GO:0002142 | 2.52E-03 |
| Canada (Newfoundland) | stereocilia ankle link | GO:0002141 | 2.52E-03 |
| Canada (Newfoundland) | stereocilia coupling link | GO:0002139 | 2.52E-03 |

**d)**

**GO:Biological process**

| Pop | term_name | term_id | adjusted_p_value |
|---|---|---|---|
| Svalbard | multicellular organismal process | GO:0032501 | 1.70E-02 |
| France (Pyrenees) | ovarian follicle development | GO:0001541 | 2.55E-02 |
| France (Pyrenees) | female gonad development | GO:0008585 | 3.42E-02 |
| France (Pyrenees) | metabolic process | GO:0008152 | 4.38E-02 |
| E.Greenland | cytotoxic T cell differentiation | GO:0045065 | 1.76E-02 |
| W.Greenland | nucleosome assembly | GO:0006334 | 1.46E-08 |
| W.Greenland | chromatin assembly | GO:0031497 | 1.31E-06 |
| W.Greenland | nucleosome organization | GO:0034728 | 7.05E-06 |
| W.Greenland | DNA packaging | GO:0006323 | 2.53E-05 |
| W.Greenland | chromatin assembly or disassembly | GO:0006333 | 2.67E-05 |
| W.Greenland | DNA conformation change | GO:0071103 | 3.95E-05 |
| W.Greenland | protein-DNA complex assembly | GO:0065004 | 6.44E-05 |
| W.Greenland | protein-DNA complex subunit organization | GO:0071824 | 2.57E-03 |
| W.Greenland | chromatin silencing | GO:0006342 | 9.17E-03 |
| W.Greenland | chromatin organization | GO:0006325 | 1.33E-02 |
| W.Greenland | homophilic cell adhesion via plasma membrane adhesion molecules | GO:0007156 | 3.78E-02 |
| W.Greenland | chromatin organization involved in negative regulation of transcription | GO:0097549 | 4.21E-02 |

| | | | |
|---|---|---|---|
| England | RNA phosphodiester bond hydrolysis, endonucleolytic | GO:0090502 | 1.13E-07 |
| England | RNA phosphodiester bond hydrolysis | GO:0090501 | 3.41E-06 |
| England | DNA integration | GO:0015074 | 3.11E-05 |
| England | nucleic acid phosphodiester bond hydrolysis | GO:0090305 | 3.43E-05 |
| Norway | eosinophil migration | GO:0072677 | 3.02E-02 |

**GO:Molecular function**

| Pop | term_name | term_id | adjusted_p_value |
|---|---|---|---|
| Iceland | RNA-DNA hybrid ribonuclease activity | GO:0004523 | 5.09E-03 |
| Iceland | endonuclease activity, active with either ribo- or deoxyribonucleic acids and producing 5'-phosphomonoesters | GO:0016893 | 1.20E-02 |
| Iceland | endoribonuclease activity, producing 5'-phosphomonoesters | GO:0016891 | 1.53E-02 |
| Iceland | endoribonuclease activity | GO:0004521 | 2.33E-02 |
| Iceland | core promoter sequence-specific DNA binding | GO:0001046 | 4.33E-02 |
| E.Greenland | molecular carrier activity | GO:0140104 | 2.69E-02 |
| E.Greenland | nuclear export signal receptor activity | GO:0005049 | 2.70E-02 |
| W.Greenland | structural constituent of cytoskeleton | GO:0005200 | 1.08E-11 |
| W.Greenland | protein heterodimerization activity | GO:0046982 | 6.64E-06 |
| W.Greenland | structural molecule activity | GO:0005198 | 1.92E-04 |
| W.Greenland | nuclear export signal receptor activity | GO:0005049 | 1.17E-02 |
| England | structural constituent of cytoskeleton | GO:0005200 | 1.78E-19 |
| England | structural molecule activity | GO:0005198 | 6.13E-11 |
| England | RNA-DNA hybrid ribonuclease activity | GO:0004523 | 6.65E-08 |
| England | endoribonuclease activity, producing 5'-phosphomonoesters | GO:0016891 | 2.51E-07 |
| England | endoribonuclease activity | GO:0004521 | 1.35E-06 |
| England | endonuclease activity, active with either ribo- or deoxyribonucleic acids and producing 5'-phosphomonoesters | GO:0016893 | 2.03E-06 |
| England | endonuclease activity | GO:0004519 | 6.03E-06 |
| England | ribonuclease activity | GO:0004540 | 1.42E-04 |
| England | RNA-directed DNA polymerase activity | GO:0003964 | 1.10E-03 |
| England | RNA stem-loop binding | GO:0035613 | 1.95E-03 |
| England | nuclease activity | GO:0004518 | 2.20E-03 |
| England | nuclear export signal receptor activity | GO:0005049 | 2.33E-03 |
| England | catalytic activity, acting on RNA | GO:0140098 | 5.69E-03 |
| England | DNA polymerase activity | GO:0034061 | 2.53E-02 |
| Canada (Newfoundland) | structural constituent of cytoskeleton | GO:0005200 | 2.53E-05 |

**GO:Cellular component**

| Pop | term_name | term_id | adjusted_p_value |
|---|---|---|---|
| Svalbard | nucleosome | GO:0000786 | 3.29E-03 |
| Svalbard | extracellular matrix | GO:0031012 | 8.96E-03 |
| France (Pyrenees) | cytoplasm | GO:0005737 | 2.43E-04 |
| France (Pyrenees) | intracellular anatomical structure | GO:0005622 | 1.86E-03 |
| France (Pyrenees) | intracellular membrane-bounded organelle | GO:0043231 | 3.13E-03 |
| France (Pyrenees) | membrane-bounded organelle | GO:0043227 | 5.57E-03 |
| Iceland | nucleosome | GO:0000786 | 1.64E-03 |
| Iceland | protein-DNA complex | GO:0032993 | 2.09E-03 |
| Iceland | chromosome | GO:0005694 | 2.40E-03 |
| Iceland | DNA packaging complex | GO:0044815 | 1.12E-02 |
| Iceland | chromatin | GO:0000785 | 4.40E-02 |
| W.Greenland | nucleosome | GO:0000786 | 8.65E-26 |
| W.Greenland | DNA packaging complex | GO:0044815 | 1.01E-24 |
| W.Greenland | protein-DNA complex | GO:0032993 | 3.72E-14 |
| W.Greenland | intermediate filament | GO:0005882 | 1.81E-11 |
| W.Greenland | intermediate filament cytoskeleton | GO:0045111 | 6.57E-10 |
| W.Greenland | chromatin | GO:0000785 | 6.13E-08 |
| W.Greenland | polymeric cytoskeletal fiber | GO:0099513 | 1.51E-04 |
| W.Greenland | chromosome | GO:0005694 | 2.05E-03 |
| W.Greenland | supramolecular fiber | GO:0099512 | 2.32E-03 |
| W.Greenland | supramolecular polymer | GO:0099081 | 2.61E-03 |
| W.Greenland | host cellular component | GO:0018995 | 3.51E-03 |
| W.Greenland | host cell | GO:0043657 | 3.51E-03 |
| W.Greenland | host intracellular region | GO:0043656 | 3.51E-03 |
| W.Greenland | host intracellular organelle | GO:0033647 | 3.51E-03 |
| W.Greenland | host intracellular membrane-bounded organelle | GO:0033648 | 3.51E-03 |
| W.Greenland | host cell part | GO:0033643 | 3.51E-03 |
| W.Greenland | host intracellular part | GO:0033646 | 3.51E-03 |
| W.Greenland | host cell nucleus | GO:0042025 | 3.51E-03 |
| W.Greenland | non-membrane-bounded organelle | GO:0043228 | 4.50E-03 |
| W.Greenland | intracellular non-membrane-bounded organelle | GO:0043232 | 4.50E-03 |
| England | intermediate filament | GO:0005882 | 3.77E-19 |
| England | intermediate filament cytoskeleton | GO:0045111 | 6.48E-19 |

| | | | |
|---|---|---|---|
| England | polymeric cytoskeletal fiber | GO:0099513 | 6.99E-09 |
| England | supramolecular complex | GO:0099080 | 9.62E-08 |
| England | supramolecular fiber | GO:0099512 | 1.14E-07 |
| England | supramolecular polymer | GO:0099081 | 1.41E-07 |
| England | MHC class II protein complex | GO:0042613 | 5.70E-03 |
| England | MHC protein complex | GO:0042611 | 1.25E-02 |
| England | cytoskeleton | GO:0005856 | 4.15E-02 |
| Canada (Newfoundland) | intermediate filament | GO:0005882 | 3.05E-04 |
| Canada (Newfoundland) | intermediate filament cytoskeleton | GO:0045111 | 4.88E-04 |
| Canada (Newfoundland) | endoplasmic reticulum subcompartment | GO:0098827 | 3.33E-02 |

**e)**

**GO:Biological process**

| Pop | term_name | term_id | adjusted_p_value |
|---|---|---|---|
| Svalbard | regulation of peptidase activity | GO:0052547 | 2.13E-03 |
| Svalbard | negative regulation of peptidase activity | GO:0010466 | 2.48E-03 |
| Svalbard | regulation of endopeptidase activity | GO:0052548 | 1.37E-02 |
| France (Pyrenees) | nucleosome assembly | GO:0006334 | 3.13E-04 |
| France (Pyrenees) | nucleosome organization | GO:0034728 | 6.38E-03 |
| France (Pyrenees) | protein-DNA complex assembly | GO:0065004 | 2.12E-02 |
| France(Alps) | cell junction disassembly | GO:0150146 | 3.75E-03 |
| Sweden | nucleosome mobilization | GO:0042766 | 4.00E-03 |
| Norway | negative regulation of peptidase activity | GO:0010466 | 1.12E-05 |
| Norway | negative regulation of hydrolase activity | GO:0051346 | 7.92E-05 |
| Norway | negative regulation of endopeptidase activity | GO:0010951 | 3.94E-04 |
| Norway | negative regulation of proteolysis | GO:0045861 | 2.79E-03 |
| Norway | negative regulation of catalytic activity | GO:0043086 | 7.51E-03 |
| Norway | regulation of peptidase activity | GO:0052547 | 1.55E-02 |

**Molecular function**

| Pop | term_name | term_id | adjusted_p_value |
|---|---|---|---|
| Svalbard | endopeptidase inhibitor activity | GO:0004866 | 1.33E-06 |
| Svalbard | endopeptidase regulator activity | GO:0061135 | 2.42E-06 |
| Svalbard | peptidase inhibitor activity | GO:0030414 | 2.79E-06 |
| Svalbard | peptidase regulator activity | GO:0061134 | 3.27E-06 |

| | | | |
|---|---|---|---|
| Svalbard | enzyme inhibitor activity | GO:0004857 | 1.35E-03 |
| Svalbard | enzyme regulator activity | GO:0030234 | 3.87E-02 |
| France (Pyrenees) | protein heterodimerization activity | GO:0046982 | 1.96E-03 |
| France (Pyrenees) | binding | GO:0005488 | 5.87E-03 |
| England | peptidase regulator activity | GO:0061134 | 3.69E-03 |
| England | endopeptidase inhibitor activity | GO:0004866 | 1.10E-02 |
| England | endopeptidase regulator activity | GO:0061135 | 1.42E-02 |
| England | peptidase inhibitor activity | GO:0030414 | 1.51E-02 |
| England | SMAD binding | GO:0046332 | 1.69E-02 |
| Canada (Newfoundland) | structural constituent of cytoskeleton | GO:0005200 | 7.40E-05 |
| Norway | endopeptidase inhibitor activity | GO:0004866 | 5.03E-09 |
| Norway | endopeptidase regulator activity | GO:0061135 | 1.02E-08 |
| Norway | peptidase inhibitor activity | GO:0030414 | 1.21E-08 |
| Norway | molecular function regulator | GO:0098772 | 1.37E-07 |
| Norway | peptidase regulator activity | GO:0061134 | 1.64E-07 |
| Norway | enzyme inhibitor activity | GO:0004857 | 2.35E-05 |
| Norway | enzyme regulator activity | GO:0030234 | 2.37E-04 |
| Norway | repressing transcription factor binding | GO:0070491 | 5.66E-03 |
| Norway | ligand-gated anion channel activity | GO:0099095 | 3.18E-02 |
| Norway | growth factor activity | GO:0008083 | 3.46E-02 |

**Cellular component**

| Pop | term_name | term_id | adjusted_p_value |
|---|---|---|---|
| France (Pyrenees) | nucleosome | GO:0000786 | 3.73E-13 |
| France (Pyrenees) | DNA packaging complex | GO:0044815 | 8.49E-12 |
| France (Pyrenees) | protein-DNA complex | GO:0032993 | 4.45E-08 |
| France (Pyrenees) | intracellular anatomical structure | GO:0005622 | 8.97E-03 |
| France (Pyrenees) | intracellular membrane-bounded organelle | GO:0043231 | 2.06E-02 |
| Iceland | mitochondrial respiratory chain complex IV | GO:0005751 | 1.98E-02 |
| Iceland | integral component of nuclear inner membrane | GO:0005639 | 4.28E-02 |
| Iceland | intrinsic component of nuclear inner membrane | GO:0031229 | 4.28E-02 |
| Iceland | respiratory chain complex IV | GO:0045277 | 4.28E-02 |
| France(Alps) | fibrinogen complex | GO:0005577 | 3.04E-02 |
| Sweden | intrinsic component of nuclear inner membrane | GO:0031229 | 5.07E-03 |
| Sweden | integral component of nuclear inner membrane | GO:0005639 | 5.07E-03 |
| Sweden | nuclear membrane protein complex | GO:0106083 | 9.49E-03 |

| | | | |
|---|---|---|---|
| Sweden | nuclear membrane microtubule tethering complex | GO:0106094 | 9.49E-03 |
| Sweden | meiotic nuclear membrane microtubule tethering complex | GO:0034993 | 9.49E-03 |
| Sweden | microtubule organizing center attachment site | GO:0034992 | 9.49E-03 |
| Sweden | NURF complex | GO:0016589 | 2.10E-02 |
| Sweden | nuclear membrane | GO:0031965 | 4.19E-02 |
| Canada (Newfoundland) | intermediate filament | GO:0005882 | 2.91E-04 |
| Canada (Newfoundland) | intermediate filament cytoskeleton | GO:0045111 | 1.47E-03 |

Table. S4. Number and fraction of missense, deleterious and loss-of-function mutations that can be found inside and outside runs of homozygosity ROH.

***Missense***

| Population | Inside ROH | Outside ROH | fraction inside | total number of mutations | total number of ROHs |
|---|---|---|---|---|---|
| **Svalbard** | 9681 | 105850 | 0.084 | 115531 | 8921 |
| **France (Alps)** | 2288 | 216132 | 0.010 | 218420 | 537 |
| **E.Greenland** | 3939 | 302111 | 0.013 | 306050 | 1663 |
| **W.Greenland** | 612 | 196448 | 0.003 | 197060 | 1239 |
| **Iceland** | 2756 | 150543 | 0.018 | 153299 | 3011 |
| **Sweden** | 786 | 101929 | 0.008 | 102715 | 389 |
| **France (Pyrenees)** | 20690 | 166226 | 0.111 | 186916 | 7466 |
| **Norway** | 0 | 289054 | 0.000 | 289054 | 564 |
| **Canada (Newfoundland)** | 4039 | 178270 | 0.022 | 182309 | 1119 |
| **England** | 0 | 232187 | 0.000 | 232187 | 1705 |

***Deleterious***

| Population | Inside ROH | Outside ROH | percentage inside | total number of mutations | total number of ROHs |
|---|---|---|---|---|---|
| **Svalbard** | 1570 | 17914 | 0.081 | 19484 | 8921 |
| **France (Alps)** | 0 | 31757 | 0.000 | 31757 | 537 |
| **E.Greenland** | 545 | 42977 | 0.013 | 43522 | 1663 |
| **W.Greenland** | 87 | 29753 | 0.003 | 29840 | 1239 |
| **Iceland** | 402 | 23198 | 0.017 | 23600 | 3011 |
| **Sweden** | 129 | 15214 | 0.008 | 15343 | 389 |
| **France (Pyrenees)** | 2941 | 26175 | 0.101 | 29116 | 7466 |
| **Norway** | 0 | 41692 | 0.000 | 41692 | 564 |
| **Canada (Newfoundland)** | 567 | 25017 | 0.022 | 25584 | 1119 |
| **England** | 0 | 36921 | 0.000 | 36921 | 1705 |

***LOF***

| Population | Inside ROH | Outside ROH | percentage inside | total number of mutations | total number of ROHs |
|---|---|---|---|---|---|
| **Svalbard** | 75 | 1282 | 0.055 | 1357 | 8921 |
| **France (Alps)** | 13 | 1656 | 0.008 | 1669 | 537 |
| **E.Greenland** | 24 | 2691 | 0.009 | 2715 | 1663 |
| **W.Greenland** | 1 | 1657 | 0.001 | 1658 | 1239 |
| **Iceland** | 16 | 1237 | 0.013 | 1253 | 3011 |
| **Sweden** | 0 | 903 | 0.000 | 903 | 389 |

| | | | | | |
|---|---|---|---|---|---|
| **France (Pyrenees)** | 154 | 1680 | 0.084 | 1834 | 7466 |
| **Norway** | 0 | 2325 | 0.000 | 2325 | 564 |
| **Canada (Newfoundland)** | 28 | 1511 | 0.018 | 1539 | 1119 |
| **England** | 0 | 2240 | 0.000 | 2240 | 1705 |

Table. S5. Models compared to estimate the influence of runs of homozygosity (ROH) and changes in effective population size ($N_e$) on population level mutation load (missense, deleterious and loss-of-function mutations), with the best fitting model in bold letters.

| **<u>Missense mutations</u>** | | **K** | **AICc** | **Delta_AICc** |
|---|---|---|---|---|
| Null | #mutations ~ 1 | 2 | -422.43 | 152.31 |
| Model1 | #mutations ~ ROH_short | 3 | -497.66 | 77.08 |
| Model2 | #mutations ~ ROH_long | 3 | -431.76 | 142.98 |
| Model3 | #mutations ~ ROH_short + ROH_long | 4 | -504.55 | 70.19 |
| Model4 | #mutations ~ SneP_harmonic_mean | 3 | -420.30 | 154.44 |
| Model5 | #mutations ~ PSMC_harmonic_mean | 3 | -564.82 | 9.92 |
| Model6 | #mutations ~ SneP_harmonic_mean + PSMC_harmonic_mean | 4 | -568.49 | 6.25 |
| Model7 | #mutations ~ ROH_short + SneP_harmonic_mean | 4 | -498.06 | 76.69 |
| Model8 | #mutations ~ ROH_long + SneP_harmonic_mean | 4 | -431.22 | 143.53 |
| ***Model9*** | ***#mutations ~ ROH_short + PSMC_harmonic_mean*** | ***4*** | ***-574.74*** | ***0*** |
| Model10 | #mutations ~ ROH_long + PSMC_harmonic_mean | 4 | -564.33 | 10.41 |
| **<u>Deleterious mutations</u>** | | | | |
| Null | #mutations ~ 1 | 2 | -652.74 | 132.83 |
| Model1 | #mutations ~ ROH_short | 3 | -734.35 | 51.22 |
| Model2 | #mutations ~ ROH_long | 3 | -658.36 | 127.21 |
| Model3 | #mutations ~ ROH_short + ROH_long | 4 | -755.19 | 30.38 |
| Model4 | #mutations ~ SneP_harmonic_mean | 3 | -650.60 | 134.97 |
| Model5 | #mutations ~ PSMC_harmonic_mean | 3 | -768.51 | 17.06 |
| Model6 | #mutations ~ SneP_harmonic_mean + PSMC_harmonic_mean | 4 | -770.71 | 14.86 |
| Model7 | #mutations ~ ROH_short + SneP_harmonic_mean | 4 | -734.71 | 50.86 |
| Model8 | #mutations ~ ROH_long + SneP_harmonic_mean | 4 | -657.15 | 128.42 |
| ***Model9*** | ***#mutations ~ ROH_short + PSMC_harmonic_mean*** | ***4*** | ***-785.57*** | ***0*** |
| Model10 | #mutations ~ ROH_long + PSMC_harmonic_mean | 4 | -766.35 | 19.22 |

| | | | | |
|---|---|---|---|---|
| **Loss-of-function mutations** | | | | |
| Null | #mutations ~ 1 | 2 | -962.64 | 69.9000 |
| Model1 | #mutations ~ ROH_short | 3 | -994.58 | 37.96 |
| Model2 | #mutations ~ ROH_long | 3 | -973.57 | 58.97 |
| Model3 | #mutations ~ ROH_short + ROH_long | 4 | -992.38 | 40.17 |
| Model4 | #mutations ~ SneP_harmonic_mean | 3 | -960.85 | 71.69 |
| Model5 | #mutations ~ PSMC_harmonic_mean | 3 | -1029.09 | 3.45 |
| ***Model6*** | ***#mutations ~ SneP_harmonic_mean + PSMC_harmonic_mean*** | 4 | ***-1032.54*** | ***0*** |
| Model7 | #mutations ~ ROH_short + SneP_harmonic_mean | 4 | -992.38 | 40.16 |
| Model8 | #mutations ~ ROH_long + SneP_harmonic_mean | 4 | -971.64 | 60.90 |
| Model9 | #mutations ~ ROH_short + PSMC_harmonic_mean | ***4*** | -1027.15 | 5.40 |
| Model10 | #mutations ~ ROH_long + PSMC_harmonic_mean | 4 | -1030.69 | 1.85 |

Table. S6. GPS coordinates of sampled individuals, depth of coverage of the sequenced individuals and presence or absence of an assembled mitochondrial genome.

| **Pop** | **ind_id** | **Long** | **Lat** | **X_coverage** | **Mt_genome** |
|---|---|---|---|---|---|
| France (Alps) | Al1 | 6.821217 | 45.561625 | 30.1453 | Present |
| France (Alps) | Al2 | 6.652426 | 46.043455 | 30.2759 | Present |
| France (Alps) | Al3 | 6.899521 | 45.930383 | 27.4934 | Present |
| France (Alps) | Al4 | 6.485849 | 45.331418 | 35.0109 | Present |
| France (Alps) | Al5 | 6.485849 | 45.331418 | 34.5835 | Present |
| France (Alps) | Al6 | 6.54638 | 45.397397 | 31.202 | Present |
| France (Alps) | Al7 | 6.821217 | 45.561625 | 36.1516 | Present |
| France (Alps) | Al8 | 6.360879 | 45.208986 | 37.8813 | Present |
| E.Greenland | GrE1 | -37.18155 | 65.5677 | 34.9734 | Present |
| E.Greenland | GrE10 | -37.18155 | 65.5677 | 34.9524 | Present |
| E.Greenland | GrE2 | -37.18155 | 65.5677 | 33.3337 | Present |
| E.Greenland | GrE3 | -37.18155 | 65.5677 | 24.3011 | Present |
| E.Greenland | GrE4 | -37.18155 | 65.5677 | 28.6793 | Present |
| E.Greenland | GrE5 | -37.18155 | 65.5677 | 37.6546 | Present |
| E.Greenland | GrE6 | -37.18155 | 65.5677 | 26.1828 | Present |
| E.Greenland | GrE7 | -37.18155 | 65.5677 | 22.4562 | Present |

| | | | | | |
|---|---|---|---|---|---|
| E.Greenland | GrE8 | -37.18155 | 65.5677 | 35.3793 | Present |
| E.Greenland | GrE9 | -37.18155 | 65.5677 | 33.3493 | **Missing** |
| Iceland | Ice1 | -16.72699 | 65.75795 | 42.67 | Present |
| Iceland | Ice10 | -17.25715 | 66.00331 | 35.6639 | Present |
| Iceland | Ice2 | -16.72975 | 65.75345 | 35.1784 | Present |
| Iceland | Ice3 | -16.78278 | 65.85958 | 28.4251 | Present |
| Iceland | Ice4 | -17.17805 | 65.62938 | 35.2815 | Present |
| Iceland | Ice5 | -17.10376 | 65.37151 | 35.269 | Present |
| Iceland | Ice6 | -16.95315 | 65.74774 | 34.6131 | Present |
| Iceland | Ice7 | -17.24829 | 66.00882 | 35.2065 | Present |
| Iceland | Ice8 | -16.47354 | 65.51888 | 31.2049 | Present |
| Iceland | Ice9 | -16.58676 | 65.5589 | 35.671 | Present |
| Sweden | JHGO938 | 11.0850549 | 62.4810141 | 31.6015 | Present |
| Sweden | JHGO939 | 11.0850549 | 62.4810141 | 35.3767 | Present |
| Sweden | JHGO940 | 11.0850549 | 62.4810141 | 30.2544 | Present |
| Sweden | JHGO941 | 11.0850549 | 62.4810141 | 27.3856 | Present |
| Sweden | JHGO942 | 11.0850549 | 62.4810141 | 35.076 | Present |
| Sweden | Jam35 | 12.986714 | 63.159248 | 27.9133 | Present |
| W.Greenland | GrW1 | -51.3156375 | 68.8230829 | 24.8187 | **Missing** |
| W.Greenland | GrW2 | -51.3156375 | 68.8230829 | 25.1484 | Present |
| W.Greenland | GrW3 | -51.3156375 | 68.8230829 | 25.3066 | Present |
| W.Greenland | GrW4 | -51.3156375 | 68.8230829 | 28.1572 | Present |
| W.Greenland | GrW5 | -51.3156375 | 68.8230829 | 26.5623 | Present |
| W.Greenland | GrW6 | -51.3156375 | 68.8230829 | 25.4617 | Present |
| W.Greenland | GrW7 | -51.3156375 | 68.8230829 | 35.2135 | Present |
| W.Greenland | GrW8 | -51.3156375 | 68.8230829 | 35.3123 | Present |
| W.Greenland | GrW9 | -51.3156375 | 68.8230829 | 34.4738 | Present |
| France (Pyrenees) | Py1 | 2.453245 | 42.548619 | 36.5385 | Present |
| France (Pyrenees) | Py10 | 2.321872 | 42.374686 | 38.8796 | Present |
| France (Pyrenees) | Py2 | 2.380971 | 42.539329 | 33.9057 | Present |
| France (Pyrenees) | Py3 | 2.108093 | 42.480257 | 39.4949 | Present |
| France (Pyrenees) | Py4 | 2.453245 | 42.548619 | 25.2753 | Present |
| France (Pyrenees) | Py5 | 2.453245 | 42.548619 | 35.3335 | Present |
| France (Pyrenees) | Py6 | 2.380971 | 42.539329 | 35.531 | Present |
| France (Pyrenees) | Py7 | 2.156962 | 42.4365 | 35.2164 | Present |
| France (Pyrenees) | Py8 | 2.156962 | 42.4365 | 35.1843 | Present |

| | | | | | |
|---|---|---|---|---|---|
| France (Pyrenees) | Py9 | 2.062733 | 42.425745 | 31.4978 | Present |
| Svalbard | Sva115 | 15.116667 | 78.016667 | 34.7996 | Present |
| Svalbard | Sva126 | 15.116667 | 78.016667 | 26.9701 | Present |
| Svalbard | Sva129 | 15.116667 | 78.016667 | 41.2309 | Present |
| Svalbard | Sva188 | 15.116667 | 78.016667 | 32.5436 | Present |
| Svalbard | SvaG294 | 15.116667 | 78.016667 | 32.2412 | Present |
| Svalbard | SvaG296 | 15.116667 | 78.016667 | 35.008 | Present |
| Svalbard | SvaG194 | 15.116667 | 78.016667 | 32.0595 | Present |
| Svalbard | Sva299 | 15.116667 | 78.016667 | 21.3322 | Present |
| Norway | Lago10 | 8.8184918 | 62.2249937 | 28.27 | Present |
| Norway | Lago11 | 8.8184918 | 62.2249937 | 27.22 | Present |
| Norway | Lago12 | 8.8184918 | 62.2249937 | 25.03 | Present |
| Norway | Lago13 | 8.8184918 | 62.2249937 | 24.13 | Present |
| Norway | Lago14 | 8.8184918 | 62.2249937 | 28.72 | Present |
| Norway | Lago15 | 8.8184918 | 62.2249937 | 25.94 | Present |
| Norway | Lago7 | 8.8184918 | 62.2249937 | 22.98 | Present |
| Norway | Lago8 | 8.8184918 | 62.2249937 | 26.71 | Present |
| Norway | Lago9 | 8.8184918 | 62.2249937 | 26.68 | Present |
| Canada (Newfoundland) | NL1001 | -55.210694 | 47.817667 | 27.0978 | Present |
| Canada (Newfoundland) | NL1003 | -55.210694 | 47.817667 | 34.9642 | Present |
| Canada (Newfoundland) | NL1007 | -55.210694 | 47.817667 | 22.4718 | Present |
| Canada (Newfoundland) | NL1010 | -58.4052375 | 47.681049 | 39.35 | Present |
| Canada (Newfoundland) | NL1011 | -58.4052375 | 47.681049 | 34.4789 | Present |
| Canada (Newfoundland) | NL1013 | -53.2265194 | 47.2902711 | 26.7116 | Present |
| England | Scot1 | -2.0640388 | 54.3788827 | 24.41 | Present |
| England | Scot2 | -2.0640388 | 54.3788827 | 24.79 | Present |
| England | Scot3 | -2.0640388 | 54.3788827 | 27.98 | Present |
| England | Scot4 | -2.0640388 | 54.3788827 | 24.314 | Present |
| England | Scot5 | -2.0640388 | 54.3788827 | 27.2 | Present |
| England | Scot6 | -2.0640388 | 54.3788827 | 24.71 | Present |
| England | Scot7 | -2.0640388 | 54.3788827 | 25.122 | Present |
| England | Scot8 | -2.0640388 | 54.3788827 | 25.34 | Present |
| England | Scot9 | -2.0640388 | 54.3788827 | 26.37 | Present |
